# Impact of Slow Wave Abnormalities and Impaired Coordination of Pyloric Closure and Antral Contraction on Gastric Emptying: A Compartmental Modeling Study

**DOI:** 10.1101/2025.11.09.687391

**Authors:** Shannon Q. Fernandes, Mayuresh V. Kothare, Roberta Sclocco, Babak Mahmoudi

## Abstract

**Purpose:** To develop a computationally efficient gastric compartmental model that simulates diseased stomach function by altering antral-pyloric coordination and “slow wave” properties. The model evaluates motility, gastric emptying and mixing. The computational efficiency of the model enables broad parameter sweeps to simulate various pathological conditions, offering an alternative to computationally expensive finite element or finite volume approaches.

**Methods:** We developed an extended compartmental model to simulate gastric function under dysrhythmic conditions. Building on prior work, this framework incorporates enhanced fluid flow equations and improved inter-compartment connectivity. These modifications enable simulation of impaired antro-pyloric coordination, regional variations in “slow wave” frequency (bradygastria, tachygastria), and reduced amplitude to mimic quiescent activity. Model outputs included gastric emptying rates, mixing efficiency, and transpyloric flow events across healthy and impaired conditions.

**Results:** Simulations demonstrated that abnormal “slow wave” frequency or amplitude delays gastric emptying and diminishes mixing efficiency. Bradygastria induced retrograde transpyloric flow, reflecting backflow from intestine to stomach, a pathological marker. These disruptions were most pronounced when antro-pyloric coordination was impaired. Predictions aligned with prior experimental and computational findings, while model execution was ***~*** 50-fold faster than real-time gastric dynamics, highlighting computational efficiency.

**Conclusion:** This physiologically inspired compartmental model captures the impact of “slow wave” abnormalities on gastric motility. By reproducing impaired flow and mixing patterns characteristic of diseased states, it provides a valuable tool for probing mechanisms of gastric dysfunction. Importantly, its computational efficiency positions the model for use in developing and rapid testing of model-based, closed-loop neurostimulation therapies for gastrointestinal disorders.

## 1 Introduction

Gastric emptying relies on stomach motility, orchestrated by rhythmic electrical “slow wave” activity from Intestitial Cells of Cajal (ICC), which translates into muscle contractions along the stomach wall [1–7]. In a healthy human stomach, these contractions, occurring at approximately 3 cycles per minute (cpm) [2, 8], originate from the gastric pacemaker region situated in the upper body of the stomach and propagate to the antrum region [8, 9]. The pyloric sphincter, located at the junction between the stomach and duodenum, regulates the flow of gastric contents into the duodenum, facilitating either gastric emptying or mixing [10–12]. Antral contractions, occurring at approximately 3 cpm, along with an open pyloric sphincter, facilitate gastric emptying. However, if the pyloric sphincter closes when antral contractions reach the terminal antrum region, a reversal of flow occurs resulting in the propulsion of gastric contents to the proximal antrum, facilitating gastric mixing [10, 13, 14].

Therefore, stomach function relies heavily on this coordinated electrical activity for accomplishing both gastric mixing and emptying. Previous studies [8, 9, 15], have shown that lack of electrical coordination or damage to cells responsible for conducting current from the neural input can lead to altered gastric flow patters, affecting emptying and mixing. Various disorders related to gastric motility include gastroesophageal reflux disease [16], bile reflux, [17, 18], dumping syndrome, cyclic vomiting syndrome [19], gastroparesis [20], pyloric stenosis [21, 22], achalasia [23, 24], and functional dyspepsia [25]. These GI disorders are caused by the impaired coordination of the periodic opening and closing behavior of the pyloric sphincter with antral contractions.

Dysrhythmias and arrhythmias in the GI system represent any abnormalities or disruptions in the “slow wave” electrical rhythm [26–29]. A characteristic of dysrhythmias is a change in frequency. Two common dysrhythmias are tachygastria and bradygastria [30, 31]. Tachygastria refers to “slow wave” activity greater than 4 cpm, while bradygastria refers to activity less than 2 cpm [32]. Some dysrhythmias involve minimal or no electrical activity, a condition known as the quiescent state [33]. Moreover, spatial dysrhythmias could also exist in the stomach, where an ectopic pacemaker initiates “slow waves” at an abnormal location in the stomach, disrupting normal conductive activity [29]. Therefore, understanding the effect of “slow wave” frequency on gastric emptying and mixing in the stomach is important for diagnosing and treating GI disorders.

However, studying the function of the stomach in its disordered state can be challenging experimentally due to limitations in experimental techniques and ethical concerns [34–36]. Alternately, focusing on computational and numerical techniques may allow researchers to understand stomach motility and gastric emptying and related disorders using a model or “digital twin” of the organ. [10, 37, 38].

To date, computational studies have primarily focused on modeling gastric electrophysiology during dysrhythmias [37, 38]. Some numerical studies have also examined the importance of antrum-pyloric sphincter coordination [10, 39]. For instance, Ishida et al., 2019 [10] studied the gastric emptying behavior during impaired coordination of the antrum-pyloric sphincter by conducting a Computational Fluid Dynamics (CFD) analysis. However, these numerical studies are computationally expensive, as the methods used are Finite Element Method (FEM) or Finite Volume Method (FVM). Such FEM/FVM models cannot be used in the rapid screening of neurostimulation therapies *in silico* using tools from optimization and control theory, where computationally inexpensive models are preferred for use in real-time [7, 40, 41]. Furthermore, none of these existing studies integrate electrophysiology, muscle mechanics, and fluid dynamics in a unified, systems-level framework, particularly under dysrhythmic conditions.

This paper aims to use a computationally efficient gastric compartmental model framework [7] to simulate the stomach function in diseased state. Our approach involves modifying the rhythmic behavior of the pyloric sphincter to emulate impaired coordination between the antrum and pyloric sphincter. Additionally, we alter the amplitude and frequency of the ICC “slow waves” to simulate conditions of bradygastria, tachygastria and quiescent states. The paper also includes derivations to model fluid backflow based on antral relaxation and delayed pyloric closure, along with modifications to compartmental fluid transport connectivity to accommodate bidirectional flow. Each simulation is evaluated based on parameters such as stomach motility, gastric emptying, and mixing efficiency, along with an assessment of computational time for each case. The simulations are also validated against prior experimental and computational findings. Since the model is computationally cheap, it allows extensive parameter sweeps to assess dysrhythmic behavior under varying conditions, an aspect that is not straightforward with computationally expensive FEM/FVM models.

## 2 Materials and Methods

### 2.1 Modeling the function of the stomach: An overview

Stomach motility is driven by rhythmic electrical activity known as “slow waves,” generated by ICC on the stomach wall and passively conducted to adjacent Smooth Muscle Cells (SMC). These slow waves trigger intracellular calcium influx, leading to tissue stress and subsequent deformations. The role of ICC “slow waves” in muscle contractions was detailed in our previous work [7].

Muscle contractions facilitate gastric liquid movement in either anterograde or retrograde directions. Anterograde flow enables gastric emptying, pushing contents into the duodenum via the Pyloric sphincter (PS), while retrograde flow promotes gastric mixing. Peristalsis in the antral region of the stomach plays a crucial role in gastric emptying [7, 10, 42]. However, when the peristaltic wave reaches the antral-pyloric (AP) or Terminal antrum (TA) region, the PS must close; otherwise, retrograde flow of duodenum fluid occurs at the PS junction [10], potentially leading to bile reflux and GI discomfort [10]. This type of retrograde flow is undesirable. In contrast, if the PS closes during peristalsis in the antrum region of the stomach, it leads to retrograde flow within the stomach which is desirable as it aids in the mixing of gastric contents [10–12].

In our previous study [7], we have modeled the complex relationship between electrical, mechanical, and fluid behavior in the stomach (see Fig. 1) using a finite number of compartments. We extend this modeling framework to account for reverse flow behavior and improved compartment connectivity, thereby allowing us to represent the disordered states of stomach function. Detailed derivations of these modeling equations is provided in the supplementary material Section 7.1

**Fig. 1:**
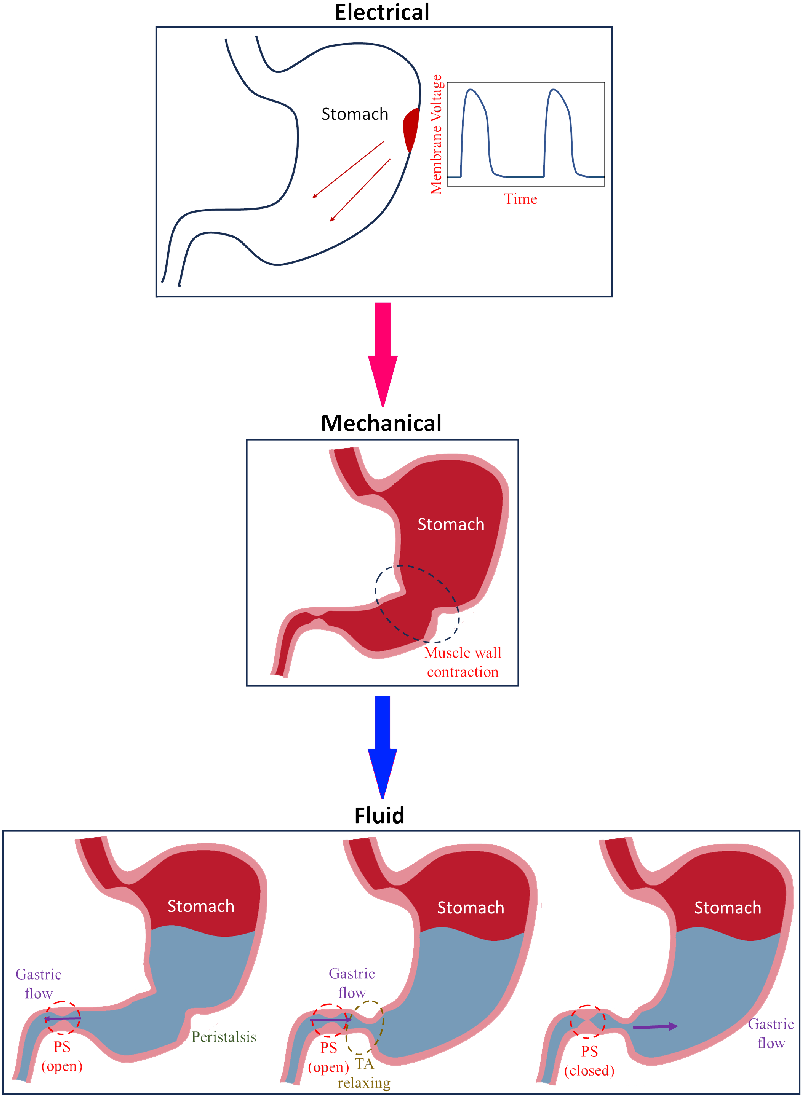
Schematic of stomach function illustrating electrical, mechanical, and fluid dynamics

### 2.2 Spatial geometry breakdown of the stomach for compartmental modeling

The spatial geometry of the stomach antrum and PS region is decomposed into four compartments (see [7]), namely, Proximal antrum (PA), Middle antrum (MA), Terminal antrum (TA), and Pyloric sphincter (PS). These regions of the stomach are chosen because they play a significant role in stomach motility during gastric emptying [13, 14]. The initial (undeformed) radii of these compartments are denoted as 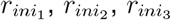, and 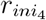, respectively, while the distances from the pacemaker region of the stomach to the compartment locations are represented by *H*_1_, *H*_2_, *H*_3_, and *H*_4_. Here, the subscript *i* represents the compartment index.

### 2.3 Modeling the electrical and mechanical behavior of the stomach

To simulate gastric electromechanics, governing equations were applied that translate ICC “slow wave” activity into muscle contractions, adapted from a previously established framework [7]. These equations (all symbols are defined in Table 2 at the end of the paper) are as follows

#### Electrical behavior

The onset of ICC “slow waves” *t*_*start*_ in a compartment is denoted as

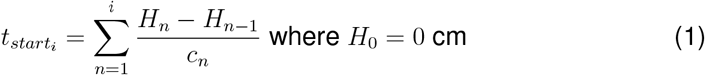

For the ICC, the “slow wave” activity is represented as

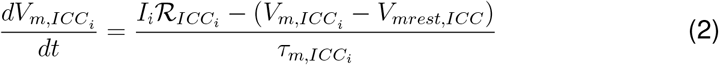

where

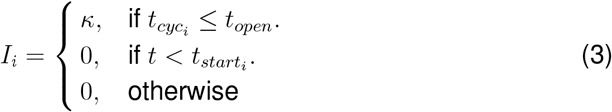

A gap junction equation connects the ICC and SMC models which is denoted by

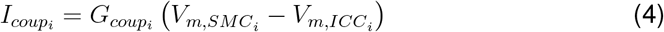

For the SMC “slow wave” activity,

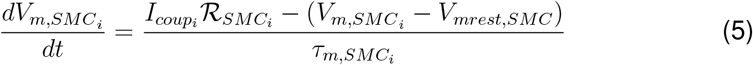

The governing equation for the intracellular concentration in the SMC for each compartment is represented as

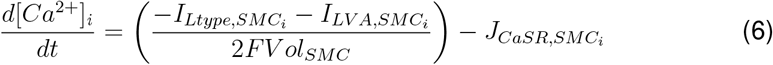

#### Mechanical behavior

The following equation represents the stress *σ* in the muscle tissue in each compartment

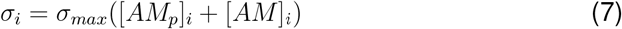

The principle stress *E* of the hyperelastic tissue model is denoted as

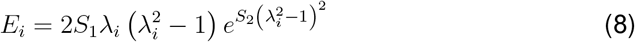

The non-linear dashpot *η*_2_ for the tissue model, which captures the hysteresis loop behavior during the loading and unloading of the material, is represented as

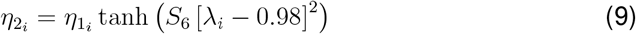

where

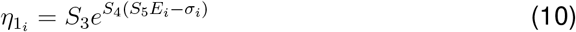

The non-linear viscoelastic model which captures the soft material tissue dynamics of the stomach tissue is denoted as

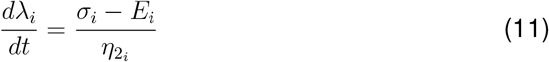

The stretched tissue strip length in each compartment is denoted as

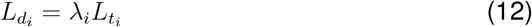

When the compartment contracts, the radius of the deformed compartment is represented as

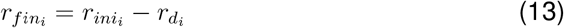

where

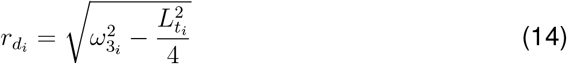

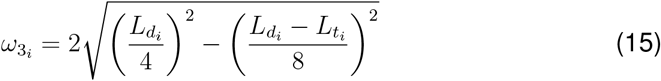

Derivation of these equations can be found in our previous paper [7].

### 2.4 Modeling the behavior of gastric liquid in the stomach

When the PS is open, gastric contents empty into the duodenum, driven by peristaltic contractions. The peristaltic flow is governed by the peristalsis equation, while the reduction in flow rate due to the PS geometry is modeled using the Bernoulli and Darcy–Weisbach equations, based on methods previously described in Fernandes et al., 2024 [7].

When the PS is closed, peristaltic contractions generate a retrograde jet, pushing gastric contents back into the antrum. This jet results from the pressure buildup between the closed PS and antral contractions and is modeled using the circular liquid jet equation, following the our previously established framework [7].

If the PS remains open during TA relaxation, retrograde flow occurs, allowing liquid from the proximal duodenum to reflux into the stomach, as observed computationally by Ishida et al., 2019 [10]. This behavior was described using the extensional flow equation.

#### 2.4.1 Modeling of peristaltic flow in gastric content transport

The velocity of gastric contents during peristaltic contractions is predicted using the peristaltic flow formulation introduced by Shapiro et al.. 1969 [43], consistent with the approach used in our earlier study [7]. The equations for the peristaltic flow in each compartment is denoted as

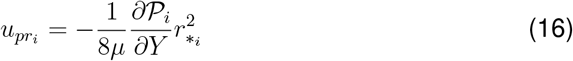

where

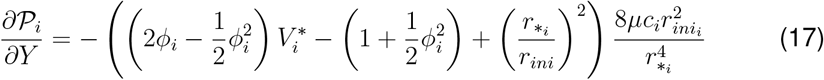

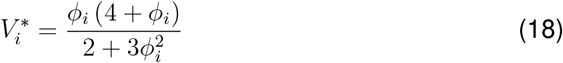

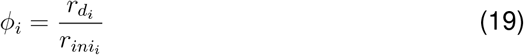

For compartments PA, MA, and TA (*i* = 1, 2, 3), the characteristic radius is defined as 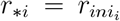 when the PS is open and 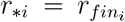 when closed. Since peristalsis does not occur in the PS, no peristaltic equation is assigned to that compartment.

The volumetric flow rate due to peristaltic contractions is given by

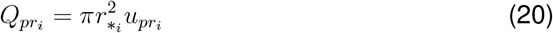

When the PS is open, peristaltic velocity 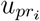 drives gastric emptying, denoted as 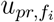, with the corresponding flow rate 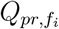. Conversely, when the PS is closed, peristaltic velocity 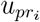 initiates a retrograde jet, represented by 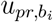.

#### 2.4.2 Retrograde jet flow initiated due to closed PS

When the PS closes, pressure is created in the region between peristaltic contraction and closed PS, which leads to a liquid jet as the gastric contents are pushed back into the antral region or body of the stomach. In our previous study [7], the liquid jet could be initiated only in the TA. However, in the present study, this restriction is removed, making the model more realistic and flexible, since the jet can initiate in any of the compartments (*i* = 1, 2, 3), provided there is peristaltic contraction and closed PS [44].

The equations governing the model of liquid jet initiation are denoted as

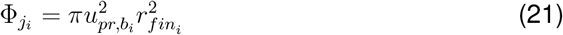

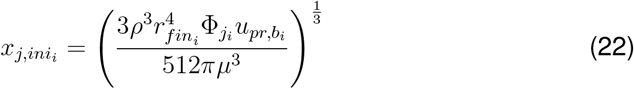

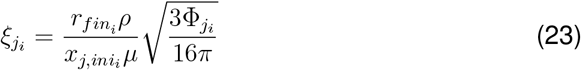

Once initiated, the liquid jet propagates to adjacent compartments [7]. Previous studies [12, 45, 46] have shown that steady-state liquid jet profiles emerge in various stomach regions when originating from the TA. In this study we simplify the liquid jet representation by focusing solely on its steady-state profile. This simplification is justified, as Ferrua and Singh, 2010 [12] demonstrated that liquid jet development occurs nearly instantaneously.

Modeling the propagation of the liquid jet depends on its point of initiation. If initiated in the TA, the jet velocity must be computed for both the MA and PA compartments (*i* = 1, 2). If initiated in the MA, velocity prediction is required only for the PA (*i* = 1). If the jet originates in the PA, no further velocity predictions are necessary. The equations to predict the jet in other compartments are represented as

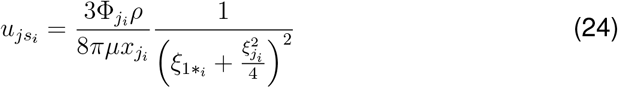

where

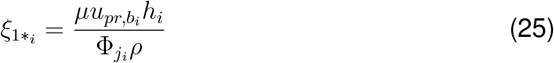

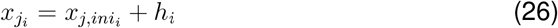

#### 2.4.3 Extensional flow in the TA

If the PS is open and the TA relaxes, retrograde flow occurs at the PS junction, causing gastric contents to flow from the intestine back into the stomach. This behavior was observed in a CFD study by Ishida et al., 2019 [10]. In this paper, the reverse flow due to TA relaxation is modeled using an extensional flow equation in the TA compartment (*i* = 3). The resulting gastric liquid velocity, *u*_*ae*_, is given by

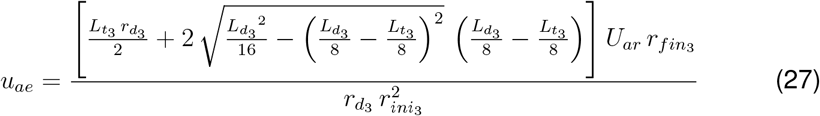

where

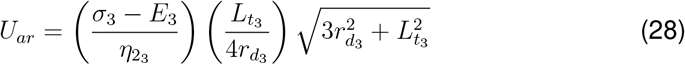

Here, the relaxation velocity of the TA compartment is denoted as *U*_*ar*_. The volumetric flow rate for the extensional flow in the TA compartment is represented as

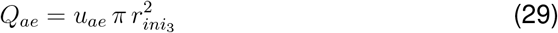

For the complete derivation of the extensional flow, see supplementary material Section 7.1.

### 2.5 Compartmental model: Interconnection to simulate continuous peristaltic flow

In the stomach, the peristaltic wave originates from the corpus region and propagates to the TA of the stomach [8, 9]. Therefore, the model must be able to predict the continuous flow of gastric liquid contents as the peristaltic wave propagates from one compartment to the next. To model this continuous behavior, differential equations are used to represent the volumetric flow rate and velocity of gastric liquid caused by peristaltic contractions when the PS is open and closed.

The equations for the continuous volumetric flow rate *Q*_*pr,f\**_ due to peristalsis when the PS is open are represented as

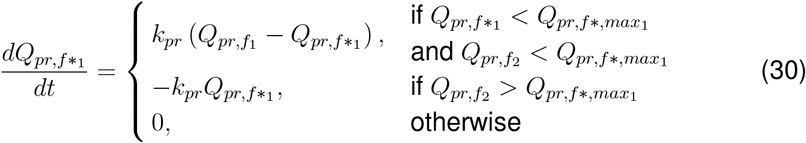

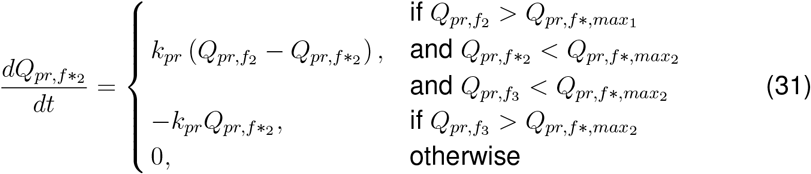

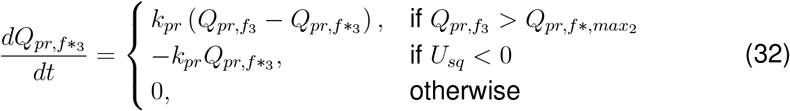

The equations for the continuous gastric liquid velocity *u*_*pr,f\**_ due to peristalsis when the PS is closed are represented as

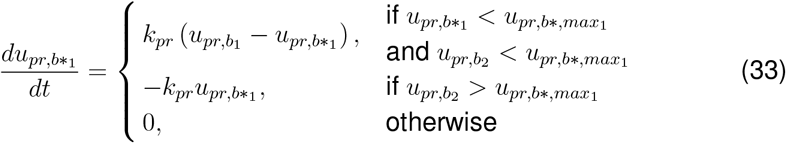

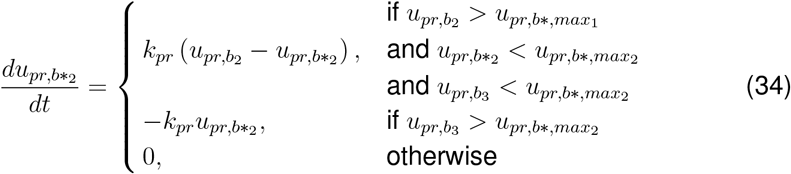

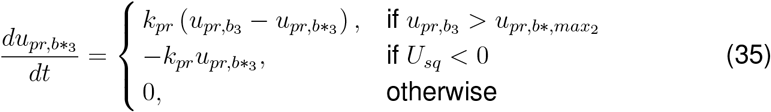

Here, *k*_*pr*_ is the rate constant that instantaneously follows the path of the peristaltic velocity. *Q*_*pr,f,max*_ represents the maximum flow rate caused by peristalsis in the given compartment when the PS is open. *u*_*pr,b,max*_ indicates the maximum velocity due to peristalsis in the given compartment when the PS is closed. For the complete derivation of this compartmental connectivity, see supplementary materials Section 7.2.

### 2.6 Discharge of gastric contents through the PS

The formulation used to describe the discharge of gastric contents through the PS is based on equations established in our previous study [7]. The velocity of gastric contents entering the PS in the direction of gastric emptying *u*_*pr\**_ is represented as

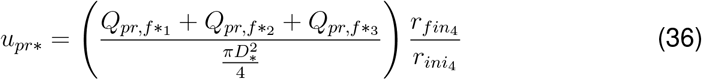

The gastric liquid emptying velocity *u*_*emp*_ equation is denoted as

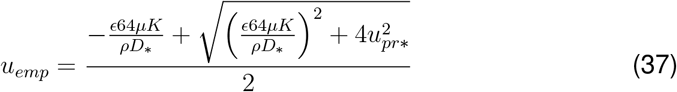

where

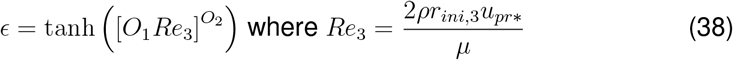

The volumetric gastric liquid emptying flow rate *Q*_*emp*_ is represented as

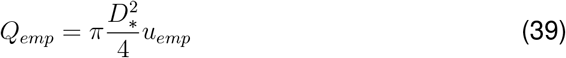

## 3 Results

The model developed in the previous sections extends our earlier formulation to incorporate a more general mechanism for reverse flow initiation and improved fluid transport dynamics. While key parameters capturing the electrophysiology, muscle mechanics, and fluid behavior of gastric contents remain consistent with our previous work [7] (also see Section 7.4.1), significant extensions have been introduced. The fluid flow component now models retrograde flow in the PS during phases of antral relaxation combined with an open sphincter. Unlike the previous model, where retrograde flow originated from the TA, the updated framework allows reverse flow to initiate dynamically from any compartment, depending on the physiological state of the antrum and PS. Furthermore, the fluid transport equations have been refined to enhance flow transport between compartments, enabling simulation of both healthy and gastric disordered states. To reflect experimental findings indicating a prolonged contraction period in the PS relative to the antrum [10, 47], several key model parameters have been modified, as described in Section 7.3.

Similar to our previous study [7], a time period variable, *T* = 20 s, represents each wave cycle, facilitating the analysis of results.

The interplay between ICC “slow wave” frequency and amplitude, controlled by *t*_*end*_ and *κ* in the model reflects the biophysical mechanisms underlying ICC pacing. Frequency modulation is governed by ion channels and intracellular signaling pathways [37, 48–50]. Prior studies have demonstrated that ICC “slow wave” frequency can be modulated by inositol 1,4,5-trisphosphate (IP3) concentration [2, 8] and calcium dynamics within pacemaker units [51]. Additionally, sodium channel impairments in ICC have been linked to reduced “slow wave” frequency, a characteristic of bradygastria [52]. Beyond frequency regulation, ion channels also influence “slow wave” amplitude. Calcium channels are critical for maintaining the plateau phase, and disruptions in their function can lead to diminished amplitude or even quiescence [2, 53]. Such disturbances of “slow wave” frequency and amplitude may contribute to gastric dysrhythmias, including tachygastria, bradygastria, and quiescence.

To simulate an unhealthy state of the stomach, the ICC “slow wave” frequency is varied in various regions by adjusting the value of *t*_*end*_. The “slow wave” frequency, in cpm, is calculated using the formula 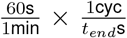. For instance, a *t*_*end*_ value of 20 seconds results in a 3 cpm frequency. Slow waves are initiated via a stimulus current *I*, where a baseline active stimulus current *κ* is applied for a healthy state. In an unhealthy state, this stimulus is altered in the antrum. For more details on parameter modifications for disease states, refer to Section 7.4.

For the gastric liquid, the base case used is a low-viscosity and low-density liquid (water), with a viscosity of 0.001 Pa.s and a density of 1000 kg*/*m^3^. For the high-viscosity gastric liquid, the density remains the same, but the viscosity is set to 10 Pa.s.

### 3.1 Comprehensive Analysis of Electrical, Mechanical, and Gastric Liquid Dynamics in the Stomach

#### 3.1.1 Healthy stomach

In a healthy stomach, ICC slow waves propagate at 3 cpm from the PA to the PS, ensuring coordinated antral and PS contractions (Fig. 2). The amplitude of 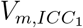 remains consistent across compartments, whereas 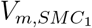 is about 31 % lower in the PA, due to a lower *G*_*coup*_ value. This reduction affects tissue stress (*σ*) and tissue stretch *λ* due to electromechanical coupling, resulting in the maximum *λ* value to be approximately 27 % lower in the PA compartment than in the other compartments.

**Fig. 2:**
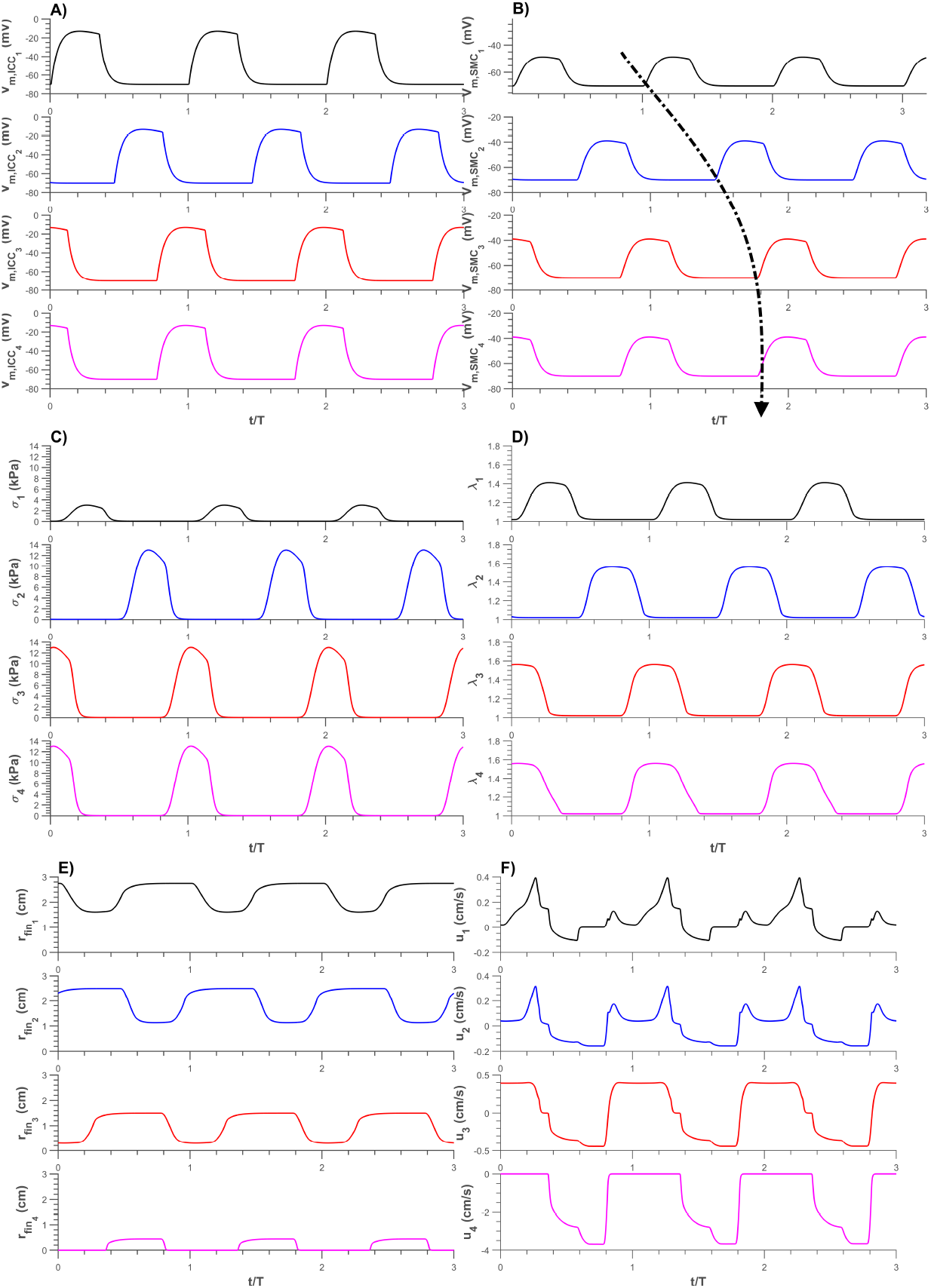
Results from the compartment model of healthy stomach (antrum + PS) at 3 cpm A) Membrane potential of ICC “slow waves” B) SMC “slow waves” C) Stress D) Stretch in muscle tissue E) Compartmental radius and F) Gastric liquid velocity in each compartment

The ICC “slow waves” drive periodic contractions, altering compartment radius *r*_*fin*_ (Fig. 2). These changes in compartmental radius drive the movement of gastric liquid. In Fig. 2 F), the negative values of liquid velocity *u*_*i*_ indicate gastric liquid movement towards the PS, which is the antegrade direction (gastric emptying direction), while the positive values indicate movement in the retrograde direction. Specifically, at the PS, the negative liquid velocity *u*_4_ indicates flow in the antegrade direction, ensuring that contents move from the stomach into the duodenum. In other compartments, both antegrade and retrograde motion occur due to retrograde jets, facilitating gastric mixing.

#### 3.1.2 Unhealthy stomach: Disrupted coordination between the antrum and the PS

For disrupted coordination, the antrum ICC “slow wave” frequency was set to 2 cpm while the PS ICC “slow wave” frequency remained at 3 cpm. The results are shown in Fig. 3. The disrupted coordination between the antrum and the PS results in the TA compartment failing to contract or relax synchronously with the PS, deviating from healthy gastric motility [7, 54]. This leads to retrograde flow velocities at the PS, as observed in Fig. 3 F).

**Fig. 3:**
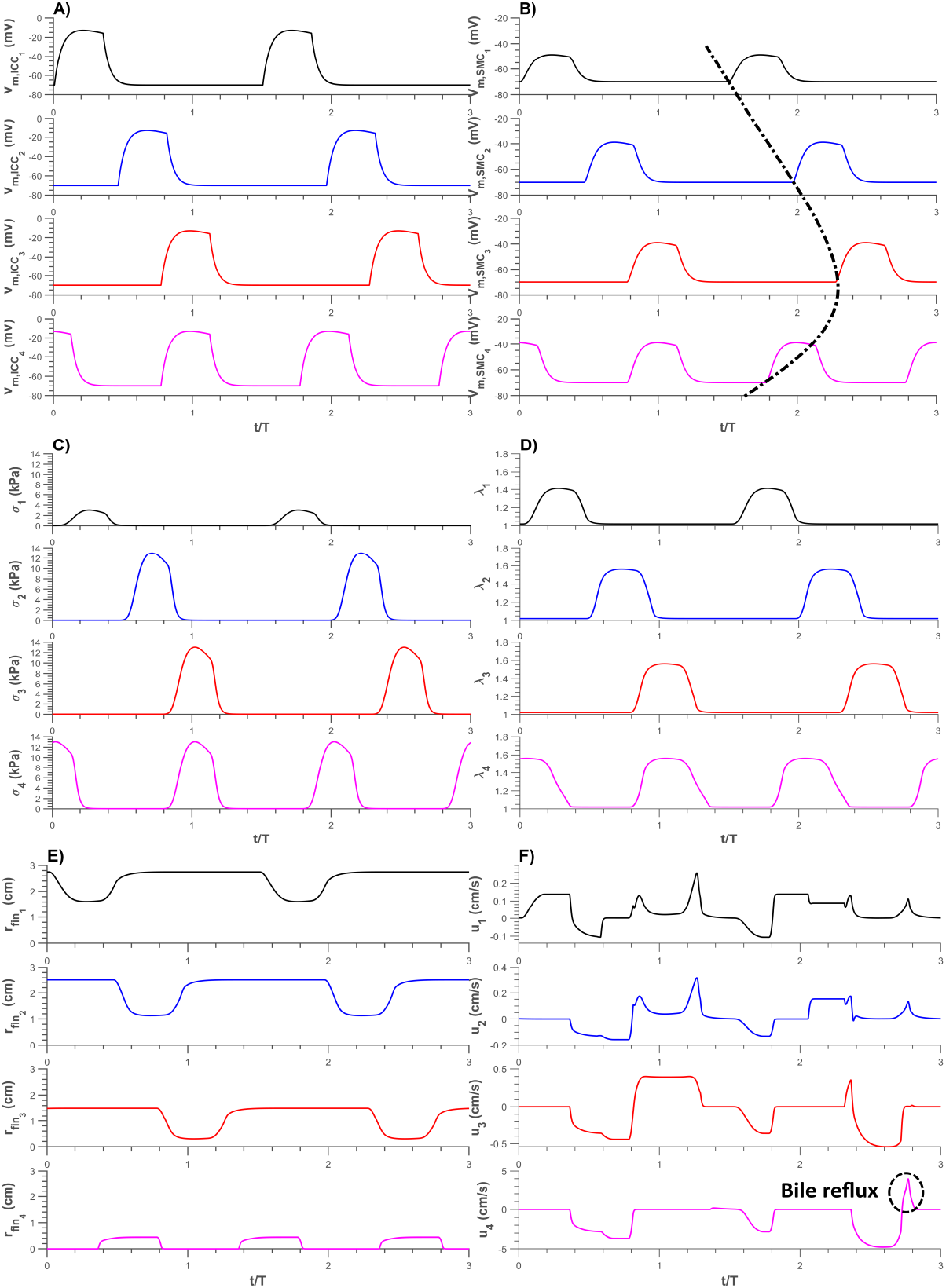
Results for a simulated unhealthy stomach, with the antrum at 2 cpm and PS at 3 cpm Membrane potential of ICC “slow waves” B) SMC “slow waves” C) Stress D) Stretch in muscle tissue E) Compartmental radius and F) Gastric liquid velocity in each compartment

#### 3.1.3 Unhealthy stomach: Change in active stimulus amplitude value

The active stimulus value *κ* was reduced to *κ*_*B*_, where *κ*_*B*_ = 0.5*κ*, for the antrum region. This reduction weakened contractions propagating from the PA compartment to the TA compartment, as shown in Fig. 4, compared to the healthy stomach case (Fig. 2). While antrum-PS coordination remained similar to that of a healthy stomach, the decrease in peristaltic amplitude significantly affected gastric emptying and mixing.

**Fig. 4:**
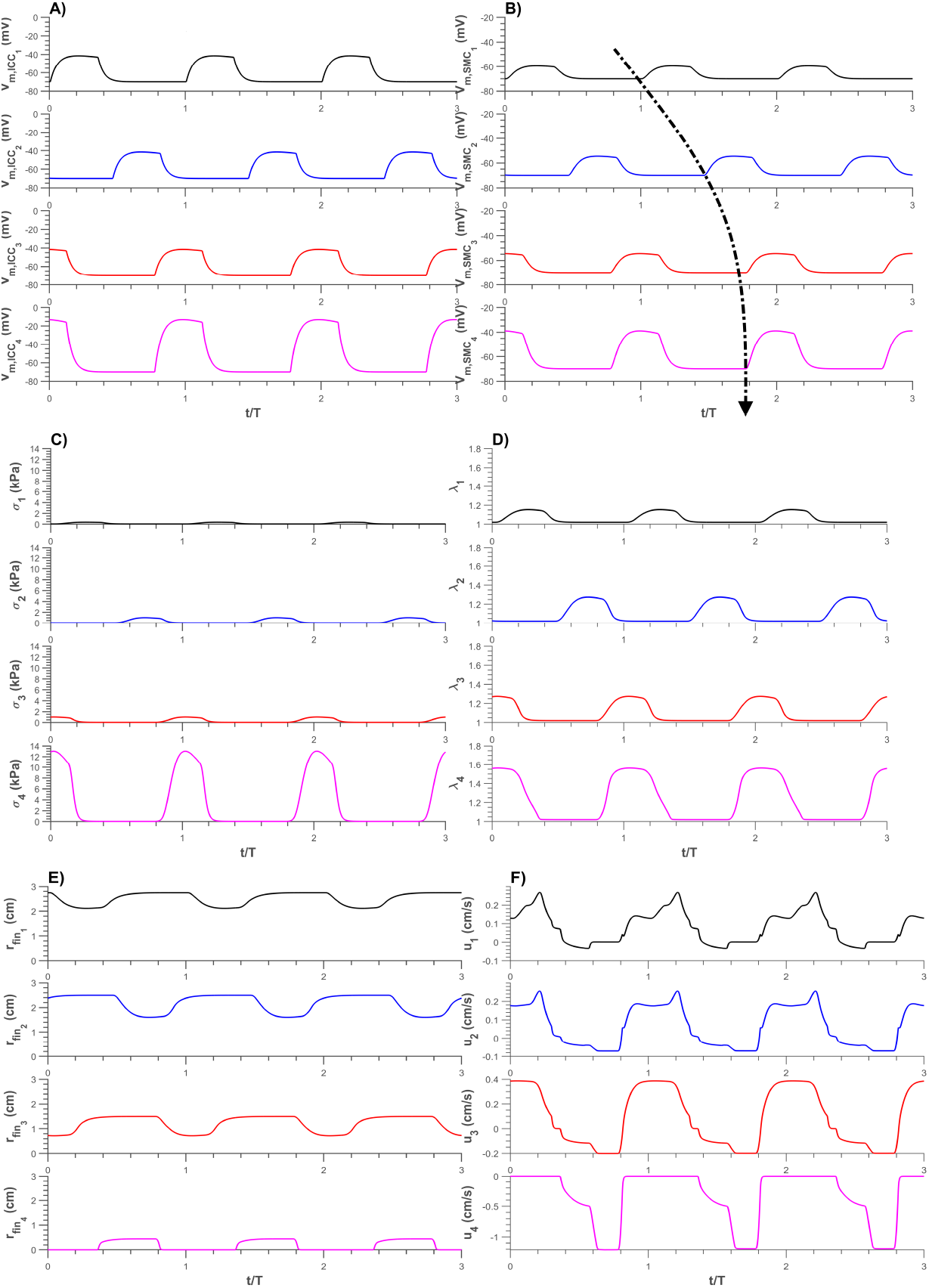
Results with the stomach slow wave at 3 cpm and reduced stimulation amplitudes for the antrum at *κ*_*B*_ and PS at *κ* A) Membrane potential of ICC “slow waves” B) SMC “slow waves” C) Stress D) Stretch in muscle tissue E) Compartmental radius and F) Gastric liquid velocity in each compartment

From Fig. 4 F), it is evident that gastric liquid flows exclusively in the antegrade direction at the PS, which is favorable for gastric emptying. However, the gastric liquid velocities are significantly lower for both gastric emptying and mixing, as seen in Fig. 4 F) compared to Fig. 2 F). Specifically, the maximum gastric liquid velocity in the PS compartment is approximately 67 % lower than in a healthy stomach.

### 3.2 Average gastric emptying rate and mixing efficiency

The method used to analyze the average gastric emptying volumetric flow rate *Q*_*emp,avg*_ and average global gastric mixing efficiency Θ_*avg*_ was based on the approach described in our previous study [7]. However, several changes and improvements were implemented in this study.

First, the average values were computed over a time period of 4 minutes, equivalent to 12 wave cycles (12 *T*), instead of using the time period of a single wave cycle. This adjustment was necessary because the current study captures disordered stomach states, including gastric liquid values with retrograde flow and impaired antrum-PS coordination. These disordered states introduce more variability into the model compared to a healthy stomach state, necessitating a longer time period for a more accurate analysis.

Moreover, after solving the model, the results were linearly interpolated using equidistant time points. This interpolation provides a more accurate and consistent method for calculating the average values.

With these improvements, the average gastric emptying volumetric flow rate *Q*_*emp,avg*_ was calculated using *Q*_*emp*_ values from 0 to 12 *T*.

The gastric mixing efficiency Θ_*i*_ in each compartment *i* was calculated using the following equation

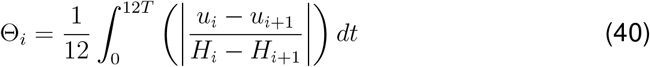

The average global gastric mixing efficiency Θ_*avg*_ is denoted as

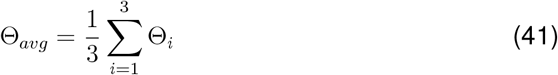

The results were computed using these analysis methods for both low viscosity and high viscosity gastric liquids.

#### 3.2.1 Healthy coordination between the antrum and PS

The ICC “slow wave” frequency of the stomach (antrum + PS) was varied while maintaining coordinated contractions between the TA and PS (Fig. 5). Specifically, PS contracted simultaneously with respect to the TA, reflecting healthy antral-PS coordination.

**Fig. 5:**
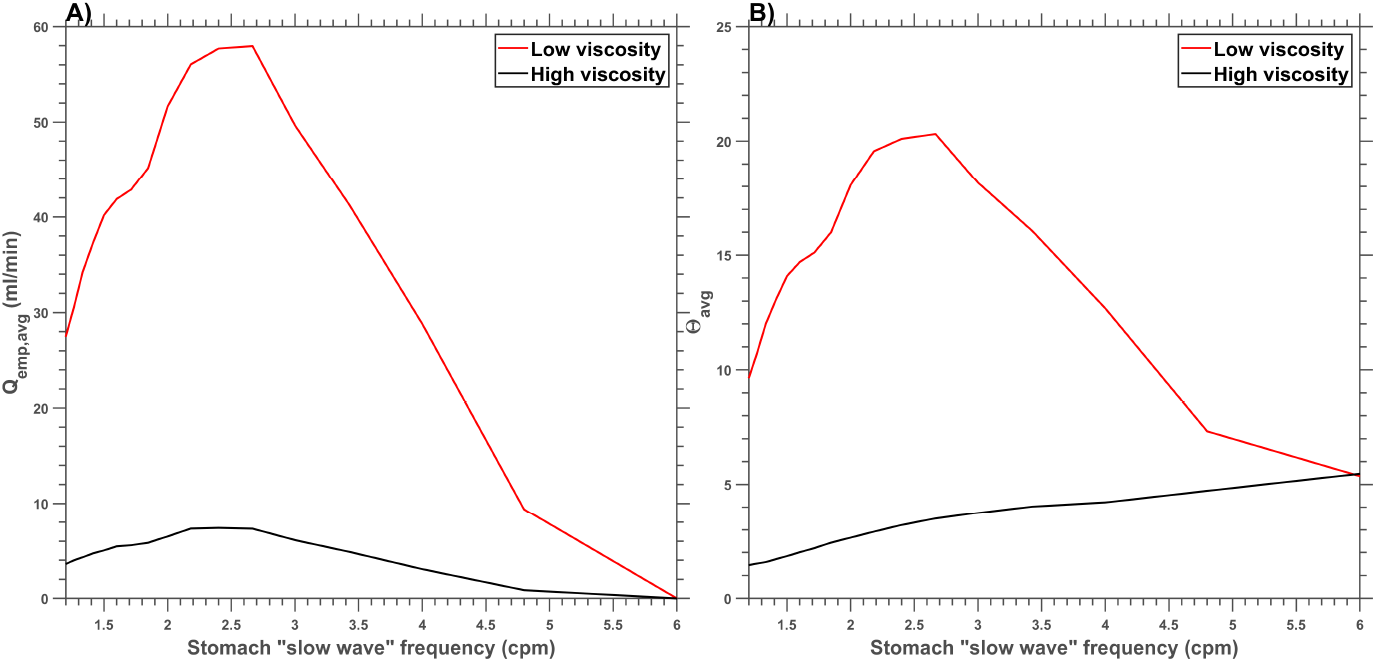
Effect of change in stomach ICC “slow wave” frequency on A) emptying rate *Q*_*emp,avg*_ B) mixing efficiency Θ_*avg*_

Previous studies have established normal “slow wave” frequencies between 2.8–3 cpm [8, 55, 56], while frequencies below 2 cpm and above 4 cpm are classified as bradygastria and tachygastria, respectively [57, 58]. Koch, 2014 [59] reported a narrower range, defining bradygastria as <2.5 cpm and tachygastria as >3.75 cpm.

Model results (Fig. 5) show that the highest *Q*_*emp,avg*_ values occurred at *~*2.7 cpm, yielding 58 ml/min and 7.3 ml/min for low- and high-viscosity gastric liquids, respectively. Corresponding Θ_*avg*_ values were 20.3 and 3.

For “slow wave” frequencies between 2.4–3 cpm, gastric emptying rate and mixing efficiency remained optimal. Below 2.4 cpm, small retrograde flows appeared at the PS, likely due to insufficient contraction duration. However, literature lacks definitive evidence explaining this phenomenon. At lower frequencies, gastric emptying and mixing efficiency declined further. Above 3 cpm, more frequent PS contractions reduced *Q*_*emp,avg*_, delaying gastric emptying.

#### 3.2.2 Impaired coordination between antrum and PS

In Fig. 6 A) and B), the ICC “slow wave” frequency for the antrum region of the stomach was maintained at 3 cpm, while the ICC “slow wave” frequency for the PS was varied. Conversely, in Fig. 6 C) and D), the ICC “slow wave” frequency for the PS was maintained at 3 cpm, while the antrum region ICC “slow wave” frequency was varied. This discrepancy in “slow wave” frequencies between the antrum and the PS leads to impaired coordination.

**Fig. 6:**
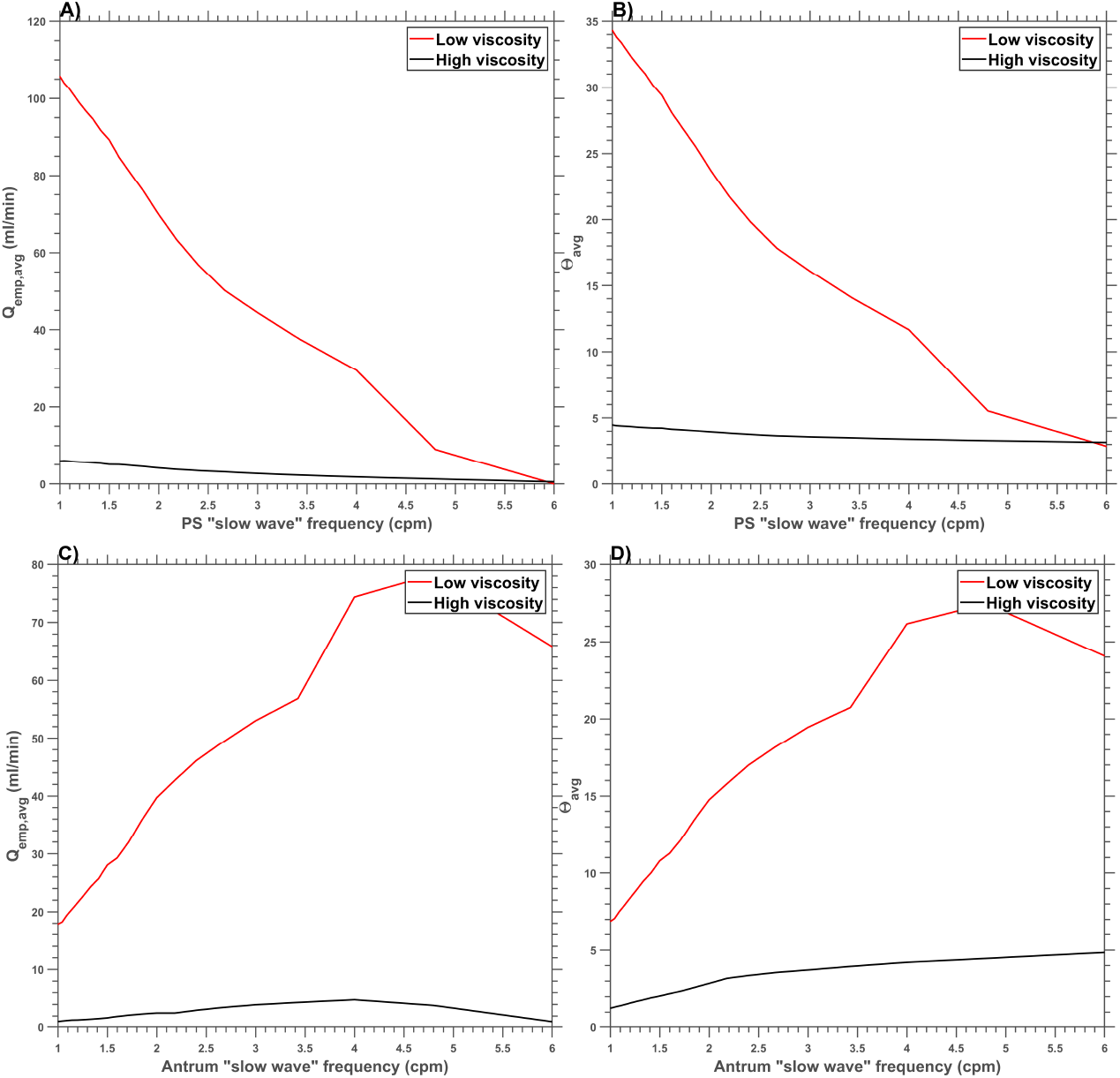
Effect of varying ICC “slow wave” frequency on gastric function A) PS frequency vs *Q*_*emp,avg*_ B) PS frequency vs Θ_*avg*_ C) Antrum frequency vs *Q*_*emp,avg*_ D) Antrum frequency vs Θ_*avg*_

In Fig. 6 A), the *Q*_*emp,avg*_ value is initially high but decreases for both high- and low-viscosity gastric liquids as the PS frequency increases. At a PS “slow wave” frequency of 6 cpm, the *Q*_*emp,avg*_ value drops to 0, indicating a persistently closed PS. For Θ_*avg*_ (Fig. 6 B)), low-viscosity liquids exhibited a steady decline with increasing PS frequency, whereas high-viscosity liquids maintained an approximately constant value of 4.7.

In Fig. 6 C) and D), *Q*_*emp,avg*_ and Θ_*avg*_ for low-viscosity liquids increased with rising antral frequency but declined beyond 4.8 cpm, likely due to insufficient relaxation time for effective peristalsis. High-viscosity liquids exhibited a similar trend, with peak *Q*_*emp,avg*_ at approximately 4 cpm. However, unlike *Q*_*emp,avg*_, Θ_*avg*_ continued to increase with increasing antral “slow wave” frequency for high-viscosity liquids.

Although the model predicts higher *Q*_*emp,avg*_ and Θ_*avg*_ at low PS frequencies (Fig. 6 A) and B)) and high antrum frequencies (Fig. 6 C) and D)), these conditions are undesirable due to retrograde flow of gastric contents into the stomach at the PS, as observed in Fig. 3.

#### 3.2.3 Change in ICC “slow wave” amplitude

In Fig. 7, the ICC “slow wave” amplitude was reduced by modifying the active stimulus current parameters to *κ*_*A*_ = 0.8*κ* and *κ*_*B*_ = 0.5*κ*. Healthy coordination between the antrum and PS was maintained while varying the “slow wave” frequency across the stomach (antrum + PS).

**Fig. 7:**
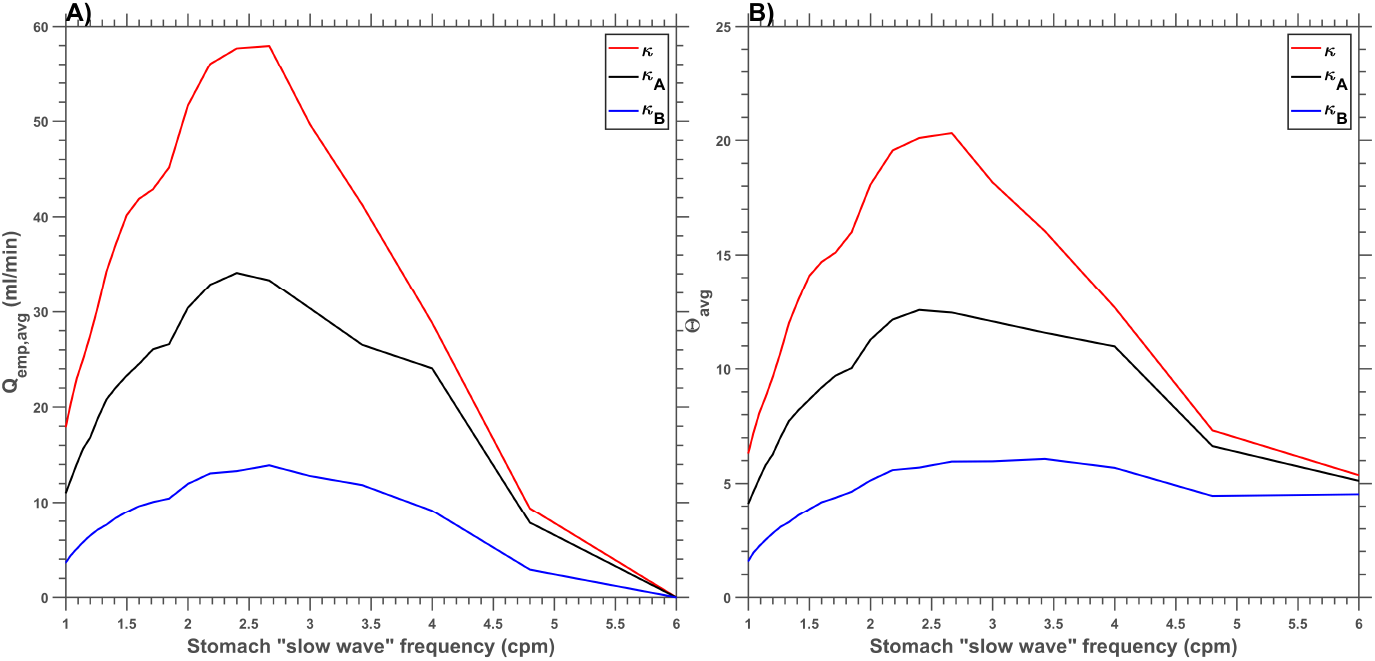
Effect of change in ICC “slow wave” amplitude on A) *Q*_*emp,avg*_ B) Θ_*avg*_

The results in Fig. 7 indicate that both the average emptying rate (*Q*_*emp,avg*_) and global mixing efficiency (Θ_*avg*_) declined with reduced ICC “slow wave” amplitude, leading to delayed gastric emptying. This effect is associated with “quiescent” slow wave activity, where minimal or no electrical activity impairs gastric motility [33].

### 3.3 Computational time

The model was solved in MATLAB utilizing various ordinary differential equation (ODE) solvers on a standard laptop featuring an Intel Core i7 7th generation processor and 16 GB of RAM. Based on our previous study [7], the healthy stomach model demonstrated optimal computational performance using the ode15s, ode23t, and ode23tb solvers. Given the structural similarity between that model and the current framework for simulating an unhealthy stomach, this study focuses on these three solvers to simulate 180 seconds of real-time gastric dynamics. The cases considered include a healthy stomach, an unhealthy stomach with a slower “slow wave” frequency for the antrum, a slower “slow wave” frequency for the PS, and a lower active stimulus value in the antrum. These cases are presented in Table 1 along with their reported computational times.

**Table 1:**
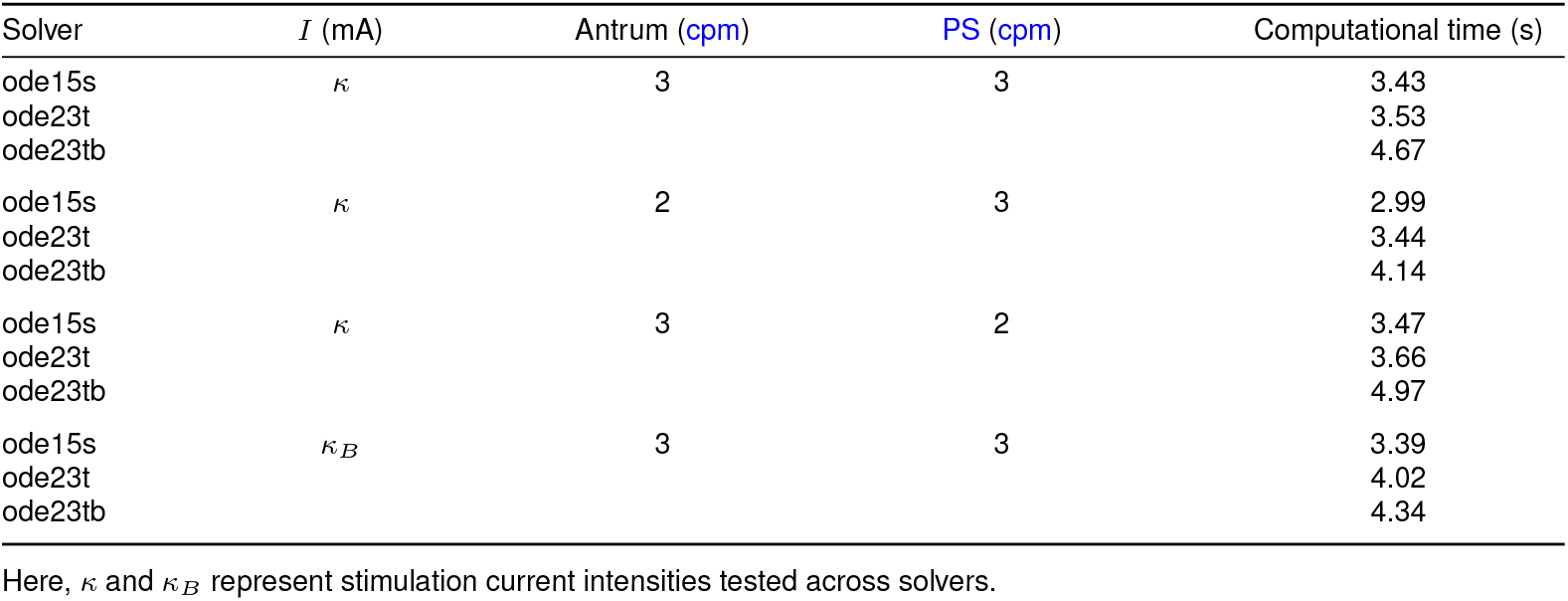
Computational time for simulating 180 s of real-time data.

From Table 1, it is evident that ode15s is the most efficient solver across all cases, consistently completing simulations in under 3.5 seconds. This performance is highly promising, as the model runs approximately 51 times faster than real-time stomach dynamics. Such computational efficiency makes the model an excellent tool for studying gastric motility using inexpensive computational resources. Additionally, the model can be employed in fields like control theory, where computationally efficient models are desirable for designing and rapid testing closed-loop control algorithms as therapies for GI disorders.

From Table 1, it is evident that ode15s is the most efficient solver across all cases, consistently completing simulations in under 3.5 seconds. This performance is highly promising, as the model runs approximately 51 times faster than real-time stomach dynamics. Such computational efficiency makes the model an excellent tool for studying gastric motility using inexpensive computational resources. Additionally, the model can be employed in fields like control theory, where computationally efficient models are desirable for designing and rapid testing closed-loop control algorithms as therapies for GI disorders.

## 4 Discussion

### Impact of “slow wave” activity on gastric function

The simulation results for the healthy stomach (Fig. 2) demonstrate reduced SMC “slow wave” amplitude in the proximal antrum and its impact on tissue stress. These findings align with previous studies that incorporate electromechanical coupling in the smooth muscle [3, 4, 6, 7]. The model also replicates physiological patterns of gastric mixing and emptying reported in the literature [6, 8, 14, 19, 54], supporting its validity in simulating healthy gastric function.

In contrast, simulations of impaired stomach conditions (Fig. 3) highlight the consequences of disrupted coordination between the antrum and PS. Increased retrograde flow at the PS suggests possible reflux of intestinal contents, which can lead to pathologies such as bile reflux [60]. These findings align with other computational studies where desynchronization between antro-pyloric compartments resulted in abnormal flow patterns like bile reflux [10], as illustrated in Fig. 8 A). Converting our simulated velocity values (Fig. 3 F) and **??** F) to instantaneous flow rates for direct comparison (note that the flow direction conventions differ: in Fig. 8 A), negative values represent bile reflux and positive values gastric emptying, whereas in our simulations the signs are reversed), we find peak bile reflux rates of *~*147 mL/min and *~* 43 mL/min, respectively. These values fall within the range reported by Ishida et al., 2019 [10], who observed a maximum reflux rate of *~*60 mL/min as shown in Fig. 8 A). Similarly, in the gastric emptying direction, our simulations show an increase in peak instantaneous flow rate from *~* 140 mL/min (Fig. 2) to *~* 190 mL/min (Fig. 3). This trend mirrors findings from Ishida et al., 2019 [10], who reported an increase in maximum emptying rate from *~* 75 to *~* 135 mL/min.

**Fig. 8:**
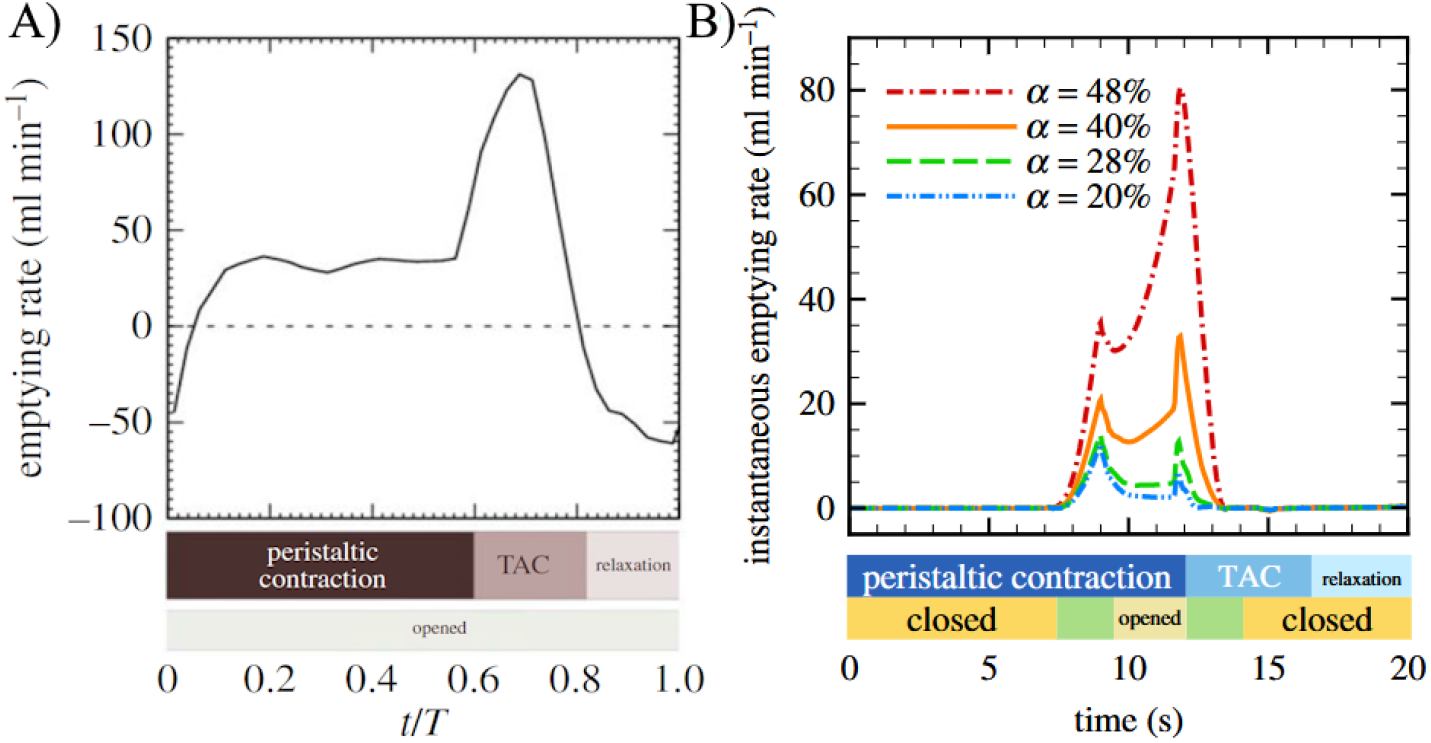
CFD studies that show A) Gastric emptying flow rate profile when the PS remains open, adapted from Ishida et al., 2019 [10]. Negative values denote bile reflux; note that our study used the opposite sign convention for our simulation results (negative for emptying, positive for reflux). This can be compared with our simulation results in Fig. 3 as described in the text. B) Gastric emptying flow rates under varying amplitudes of antral peristaltic contraction percentages (*α*), obtained from a study by Ebara et al., 2023 [62], for comparison with our simulation result in Fig. 4 as described in the text.

Additionally, from our simulation results, increasing the PS “slow wave” frequency leads to delayed emptying, and at 6 cpm, the model predicts a fully closed PS with no transpyloric flow as shown in Fig. 6. This may be due to phase delays between “slow waves” and mechanical contraction, as previously described in muscle response studies [3–5].

Reductions in either the active stimulus amplitude or ICC generated “slow wave” amplitude lead to weaker peristalsis and slower fluid velocities, consistent with delayed gastric emptying observed in motility disorders [20, 37, 61]. The simulation result shown in Fig. 4 aligns with the work of Ebara et al., 2023 [62] (Note that the sign convention is reversed in our simulation results: in Fig. 8 B), positive values represent gastric emptying, whereas in our simulations gastric emptying is represented by negative values), who showed that reduced contraction amplitude directly impairs gastric transport and emphasized maintaining normal contraction ratios (such as an 80 % contraction ratio in the distal antrum) for effective gastric emptying, as shown in Fig. 8 B). For instance, when the parameter *κ* in our model was reduced by half, impairing the strength of gastric contractions, the maximum instantaneous gastric emptying flow rate decreased to *~* 47 mL/min (when calculated from Fig. 4), compared to *~* 135 mL/min in the healthy case (Fig. 2 F)). This reduction closely matches the trend reported by Ebara et al., 2023, where lowering the contraction percentage (*α*) led to reduced instantaneous gastric flow rates.

Together, these results underscore the importance of coordinated electrical activity between the antrum and PS in supporting normal gastric function. Disruptions to amplitude or timing of “slow waves”, whether intrinsic or extrinsic, can have substantial effects on motility, highlighting key mechanisms underlying gastric dysfunction.

### Stochastic equation to simulate “slow wave” abnormalities

Until now, this study has analyzed each “slow wave” abnormality independently and in a deterministic setting. However, real-life gastric diseases often involve a combination of these abnormalities possibly appearing stochastically, as highlighted in previous studies [63, 64]. To model such mixed “slow wave” abnormalities, our study can be modified to allow the key variables, namely, “slow wave” frequency and amplitude, to vary stochastically. For instance, the stochastic variation can be described by a Stochastic Differential Equation (SDE) based on geometric Brownian motion [65], discretized using the Euler-Maruyama method, as follows:

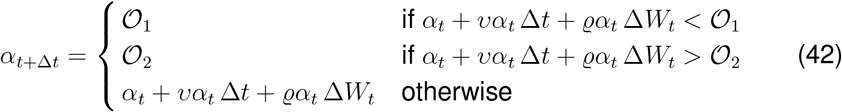

Here, *α*_*t*_ represents the value of the variable (either slow wave frequency or amplitude) at time *t, υ* denotes the average rate of change, *ϱ* represents the standard deviation of random fluctuations, Δ*t* is the time step, and Δ*W*_*t*_ is the Wiener process increment, represented as 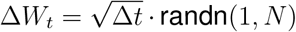, where *N* is the number of time steps. By including a lower bound *𝒪*_1_ and an upper bound *𝒪*_2_, the equation ensures that *α*_*t*_ stays within the range [*𝒪*_1_, *𝒪*_2_].

This equation can be applied to simulate stochastic variations in “slow wave” frequency or amplitude of active stimulus current. For instance, bradygastria can be modeled by setting *𝒪*_1_ = 1.8 and *𝒪*_2_ = 2.3, while tuning *υ* and *ϱ* to control fluctuation dynamics. A separate SDE can similarly govern fluctuations in the active stimulus current (*κ*). Using these equations, the effect of dynamic variations in both variables on motility and emptying can be studied, enabling the simulation of more realistic gastric dysrhythmias.

However, the scope of this study remains focused on analyzing each “slow wave” abnormality in isolation. This approach allows for a more in-depth study of the dysrhythmias associated with disordered stomach states. A future study will focus on investigating the effects described above.

### Comparative Analysis of Compartmental Stomach Model Predictions with Clinical Outcomes in Gastric Disorder

Gastric slow wave frequency dysrhythmias, such as bradygastria and tachygastria, have been linked to various gastric disorders, including functional dyspepsia (FD), gastro-esophageal reflux disease (GERD), and gastroparesis/nausea and vomiting syndromes (NVS) [66, 67]. The literature suggests that these dysrhythmias are most prominent in NVS and GERD, followed by FD. Common symptoms associated with these conditions include abdominal pain, nausea, vomiting, early satiety, postprandial fullness, bloating, and gastric discomfort [68–70]. Research studies have shown that disorders such as gastroparesis and FD are often characterized by delayed gastric emptying, which can be attributed to bradygastria, tachygastria, or reduced antral contractile amplitude [61, 68, 71–76].

The simulation results from this study, as demonstrated in Fig. 5, show that both high and low slow wave frequencies (bradygastria and tachygastria) are associated with delayed gastric emptying. Specifically, the average gastric content flow rate was lower at these frequencies compared to normogastria (approximately 2.8-3.0 cpm). An important finding is that when the PS phasic contractions are synchronized with antral contraction frequency, tachygastria can result in delayed gastric emptying. This occurs because the PS remains predominantly in its closed state, significantly reducing the gastric emptying rate. This mechanism, not previously reported in the literature, may explain the delayed gastric emptying observed in FD patients with tachygastria [72–74].

Earlier studies, such as those by Pfaffenbach et al, 1997 [75] showed that tachygastria leads to delayed gastric emptying. Ebara et al al., 2023 [62] proposed that this phenomenon may occur due to low antral contractile amplitude or impaired antral-PS coordination, though the exact mechanism remained unclear. Our findings provide new insights, suggesting that PS closure during elevated slow wave frequencies could be a key factor contributing to delayed gastric emptying as shown in Fig. 5 and 6.

Conversely, a study by Wang et al., 2021 [77] has reported that tachygastria is sometimes associated with rapid gastric emptying, along with unexplained symptoms such as chronic nausea. Our simulations (Fig. 6) provide an explanation for this phenomenon by showing that when the antral slow wave frequency increases but the PS frequency remains constant, the stomach may empty more rapidly. This impaired coordination between the antrum and PS can also lead to bile reflux, which could explain the symptoms of nausea or vomiting, as illustrated in Fig. 3 F).

Additional studies, such as those by Meier et al., 2002 [78] on patients with type 1 diabetes, Rochira et al., 2022 [68] on gastroparesis patients, and Sclocco et al., 2022 [61] on FD patients, reported that reduced peristaltic amplitude could lead to delayed gastric emptying. These findings are consistent with our results (Fig. 4 and 7), which demonstrate that lower slow wave amplitude in the antrum results in delayed gastric emptying compared to a healthy stomach.

Another significant finding from this simulation study is that at very low PS frequencies, where the PS remains predominantly open, even during antral relaxation (as shown in Fig. 3 and 6), rapid gastric emptying and bile reflux can occur. This condition is associated with dumping syndrome, as reported by Ishida et al., 2019 [10]. Prolonged bile exposure in the stomach, being an irritant, may lead to intestinal metaplasia and increase the risk of progressing to gastric cancer [60].

**In conclusion**, the computational model developed effectively simulates gastric function under healthy and pathological states, showing that optimal emptying and mixing require antral-pyloric coordination and a “slow wave” frequency of 2.4 to 3 cpm. Deviations in TA or PS frequencies reduce gastric efficiency and can induce bile reflux, while reductions in “slow wave” amplitude also impair motility. The model achieves these predictions with high computational efficiency, simulating 180 seconds of dynamics in under 3.5 seconds. Although it currently omits factors such as gastric friction, fundus and pyloric tonic contractions, neural inputs, and chemical regulation, its adaptable framework allows for future inclusion of these mechanisms to enhance physiological accuracy.

## 5 Declarations

### 5.1 Ethics approval and consent to participate

Not applicable since this is a computational study and no human or animal subjects were involved.

### 5.2 Consent for publication

All authors have read and approved the final manuscript and consent to its publication.

### 5.3 Availability of data and material

The simulation code associated with this paper is available on GitHub at: https://github.com/shanferns/Stomach-compartmental-model.git

### 5.4 Competing interests

The authors declare that they have no competing interests.

### 5.5 Funding

This work was supported in part by the National Institutes of Health, USA, Grant OT2OD030535 under the Stimulating Peripheral Activity to Reduce Conditions (SPARC) program; the John C. Chen graduate fellowship from Chemical and Biomolecular Engineering at Lehigh University for Mr. S. Fernandes; and a Faculty Innovation Grant (FIG) from Lehigh University.

### 5.6 Authors’ contributions

Shannon Q. Fernandes: Writing – review & editing, Writing – original draft, Validation, Methodology, Investigation, Formal analysis, Conceptualization. Mayuresh V. Kothare: Writing – review & editing, Writing – original draft, Validation, Supervision, Resources, Project administration, Methodology, Investigation, Funding acquisition, Formal analysis, Conceptualization. Roberta Sclocco: Writing – review & editing, Supervision, Resources, Project administration, Investigation, Conceptualization. Babak Mahmoudi: Writing – review & editing, Supervision, Resources, Project administration, Investigation, Conceptualization.

## 5.7 Acknowledgements

The authors acknowledge National Institutes of Health (NIH) and Lehigh University for their support and funding.

## 6 Nomenclature

**Table 2:** Notations.

| Notation | Description |
| --- | --- |
| $t_{start}$ | Onset of ICC “slow wave” activity |
| $t_{cyc}$ | “Slow wave” cyclic time |
| $t_{open}$ | Time for which the influx ion channels remain open |
| $c$ | Velocity for “slow wave” propagation |
| $V_{m,ICC}$ | Membrane potential of ICC |
| $V_{m,SMC}$ | Membrane potential of SMC |
| $V_{mrest,ICC}$ | Resting membrane potential of ICC |
| $V_{mrest,SMC}$ | Resting membrane potential of SMC |
| $\mathcal{R}_{ICC}$ | Resistor component of ICC |
| $\mathcal{R}_{SMC}$ | Resistor component of SMC |
| $\tau_{m,ICC}$ | Time-varying variable of ICC cell membrane |
| $\tau_{m,SMC}$ | Time-varying variable of SMC cell membrane |
| $G_{coup}$ | Gap channel coupling conductance |
| $I_{coup}$ | Gap channel coupling current |
| $I$ | Stimulus current in the ICC |
| $\kappa$ | Active stimulus current |
| $[Ca^{2+}]$ | SMC intracellular calcium concentration |
| $I_{Ltype,SMC}$ | SMC L-type calcium channel current |
| $I_{LVA,SMC}$ | SMC low-voltage activated calcium channel current |
| $J_{CaSR,SMC}$ | SMC sarcoplasmic reticulum calcium pump flux |
| $F$ | Faraday’s constant |
| $Vol_{SMC}$ | Volume of SMC cell |
| $[AM]_p$ | Total cross-bridges (actin–phosphorylated myosin) |
| $[AM]$ | Total cross-bridges (actin–myosin) |
| $\sigma_{max}$ | SMC maximum achievable active stress |
| $S_{1/2/3/4/5/6}$ | Muscle tissue constants |
| $\lambda$ | Muscle tissue stretch |
| $L_t$ | Initial length of the tissue strip |
| $r_d$ | Radius of deformation |
| $\phi$ | Peristaltic wave amplitude ratio |
| $V^*$ | Dimensionless time-mean volumetric flow rate |
| $\mu$ | Gastric liquid viscosity |
| $\rho$ | Gastric liquid density |
| $\Phi_j$ | Kinetic momentum of liquid jet |
| $\xi_j$ | Dimensionless thrust constant of liquid jet |
| $x_{j,ini}$ | Reference point distance of liquid jet |
| $\xi_{1*}$ | Dimensionless constant for thrust region area |
| $h$ | Distance to predict the liquid jet velocity |
| $u$ | Gastric liquid velocity in each compartment |
| $D_*$ | Average diameter of TA and proximal duodenum |
| $K$ | Friction loss constant |
| $\epsilon$ | Friction loss correction factor |

**Table 2 – continued from previous page**
| Notation | Description |
| --- | --- |
| $Re_3$ | Reynolds number in TA |
| $O_{1/2}$ | Friction loss correction factor constants |

## Notes

### Competing Interest Statement

The authors have declared no competing interest.

