## Supplemental material for "Impact of Slow Wave Abnormalities and Impaired Coordination of Pyloric Closure and Antral Contraction on Gastric Emptying: A Compartmental Modeling Study"

### 7 Supplementary Material: Details of Model Development for the manuscript "Impact of Slow Wave Abnormalities and Impaired Coordination of Pyloric Closure and Antral Contraction on Gastric Emptying: A Compartmental Modeling Study", by S. Q. Fernandes, M. V. Kothare, R. Sclocco, B. Mahmoudi. MATLAB model files with documentation can be found on Github at the following link https://github.com/shanferns/Stomach-compartmental-model.git

#### 7.1 Axis-symmetry extension flow

Squeeze flow occurs when two parallel plates move towards each other, compressing the material between them [79]. Conversely, when these parallel plates move apart, the resulting flow is referred to as extensional flow in this study. In the stomach, extensional flow describes the movement of gastric contents during the relaxation of the TA. This reverse flow specifically occurs at the junction of the PS when the PS is open and the TA relaxes, as demonstrated by Ishida et al. [10]. Therefore, it is necessary to derive an equation to explain this phenomenon.

To derive the extensional flow, a method similar to the squeeze flow derivation presented by Deen, 1998 [79] is used. For axial symmetry extension flow in a pipe, the fluid velocities in the axial and radial directions are denoted by $u_a$ and $u_{ar}$, respectively. The pressure drop in the axial direction is represented by $\frac{\partial \mathcal{P}}{\partial Y}$, and the velocity at which the plates (in this case, the TA tissue) open to stretch the gastric liquid flow is denoted by $U_{ar}$ as illustrated in Fig. 9.

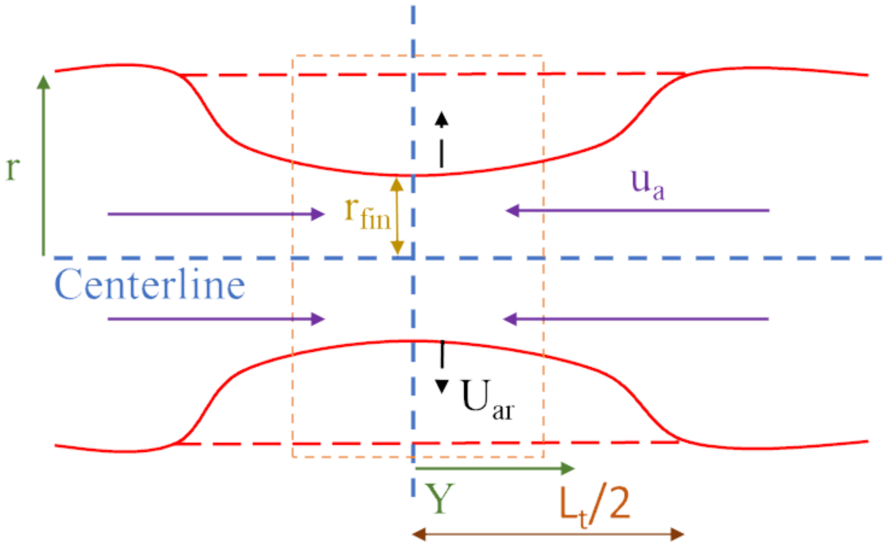


**Fig. 9**: Schematic diagram of axis symmetry extensional flow

Using the assumptions made by Deen, 1998 [79], the equation for extensional flow is considered to be pseudo-steady and almost unidirectional. Therefore, the Navier-Stokes equation is written as

$$0 = -\frac{\partial \mathcal{P}}{\partial Y} + \mu \frac{1}{r}\frac{\partial}{\partial r}\left(r\frac{\partial u_a}{\partial r}\right) \tag{S.1}$$

Boundary conditions

$$u_a(0) \text{ is bounded}$$
$$u_a(r_{fin}) = 0$$

Integrating equation S.1 and applying the boundary conditions leads to

$$u_a(r, Y, t) = \frac{-1}{4\mu}\frac{\partial \mathcal{P}}{\partial Y}\left[r_{fin}^2 - r^2\right] \tag{S.2}$$

Using continuity

$$\frac{1}{r}\frac{\partial}{\partial r}(ru_{ar}) + \frac{\partial u_a}{\partial Y} = 0 \tag{S.3}$$

Integrating equation S.3 leads to an equation for $u_{ar}$.

$$ru_{ar} = -\int_0^R r\frac{\partial u_a}{\partial Y}dr \tag{S.4}$$

The pressure gradient is dependent on the length of the tissue $L_{ta} = \frac{L_t}{2}$ and is denoted by

$$\frac{\partial \mathcal{P}}{\partial Y} = L_{ta}f(t) \tag{S.5}$$

By using equations S.2, S.4 and S.5, and a boundary condition to evaluate the integral for $u_{ar}$ which can be written as $u_{ar}(r_{fin}) = -U_{ar}$, the pressure gradient is denoted by

$$\frac{\partial \mathcal{P}}{\partial Y} = \frac{-16\mu U_{ar}L_{ta}}{r_{fin}^3} \tag{S.6}$$

The pressure gradient equation is then inserted in equation S.2, and the fluid velocity for the axis symmetry extension flow is denoted by

$$u_a = \frac{2U_{ar}L_t}{r_{fin}^3}\left[r_{fin}^2 - r^2\right] \tag{S.7}$$

To calculate $U_{ar}$ in Eq. S.7, the deformed radius of the compartment $r_d$ is used, as this radius value is dependent on the stretch of the tissue and the value of $r_d$ changes when the compartment contracts and relaxes.

To begin the derivation of $U_{ar}$, the equation for $r_d$ is expressed in terms of $\lambda$ and is denoted as

$$r_d = \sqrt{\frac{{L_t}^2\,\lambda^2}{4} - \frac{{L_t}^2}{4} - 4\left(\frac{L_t}{8} - \frac{L_t\,\lambda)}{8}\right)^2} \tag{S.8}$$

Expressing Eq. S.8 in terms of $\lambda$ when the value of $\lambda \geq 1$ can be represented as

$$\lambda = -\frac{L_t - 8\sqrt{\frac{{L_t}^2}{4} + \frac{3\,{r_d}^2}{4}}}{3\,L_t} \tag{S.9}$$

Taking the derivative of Eq. S.9 w.r.t time

$$\frac{d\lambda}{dt} = \frac{2r_d}{L_t\sqrt{\frac{3r_d^2}{4} + \frac{L_t^2}{4}}} \frac{dr_d}{dt} \tag{S.10}$$

In Eq. S.10, the derivative of $r_d$ w.r.t time represents the velocity at which the tissue walls contract or relax, thus $\frac{dr_d}{dt} = U_{ar}$. From Eq. 11, $\frac{d\lambda}{dt} = \frac{\sigma - E}{\eta_2}$ which results S.10 to be written as

$$U_{ar} = \left(\frac{\sigma - E}{\eta_2}\right)\left(\frac{L_t}{4r_d}\right)\sqrt{3r_d^2 + L_t^2} \tag{S.11}$$

The equation derived for squeeze/extensional flow is verified using a CFD software called COMSOL. The packages used in COMSOL for this simulation include the laminar flow and deforming mesh modules. The geometry chosen for the COMSOL simulation, which is in cylindrical coordinates, is shown in Fig. 10.

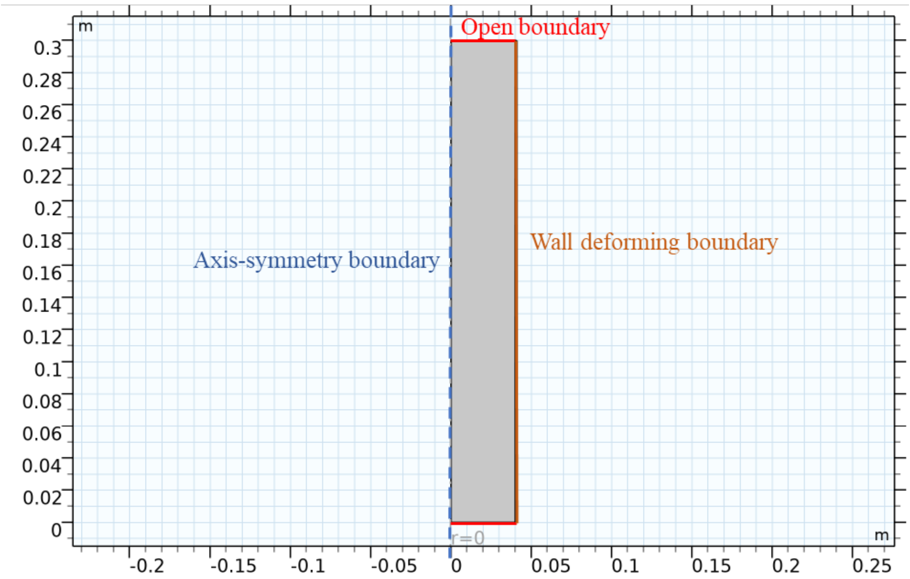


**Fig. 10**: Geometry for squeeze/extensional flow simulation in COMSOL

For the wall-deforming boundary $r_{deformed}$, a hyperbolic tangent function that is dependent on the axial length $Y$ is used, which has the form

$$r_{deformed} = A_1 \tanh\left(A_2\left(Y - A_3\right)\right) - A_1 \tag{S.12}$$

The amplitude $A_1$ of the tanh function is denoted as

$$A_1 = \begin{cases} 0 & 0 \le t \le 0.1 \\ 0.015 & 3 \le t \le 6 \\ 0 & 9 \le t \le 10 \end{cases} \tag{S.13}$$

A piecewise cubic interpolation is used, where the slope increases for $0.1 < t < 3$ and decreases for $6 < t < 9$.

The values for $A_2$ and $A_3$ are given as $250\pi$ and $0.15$.

The velocity vectors of the extensional and squeeze flow are shown in Fig. 11.

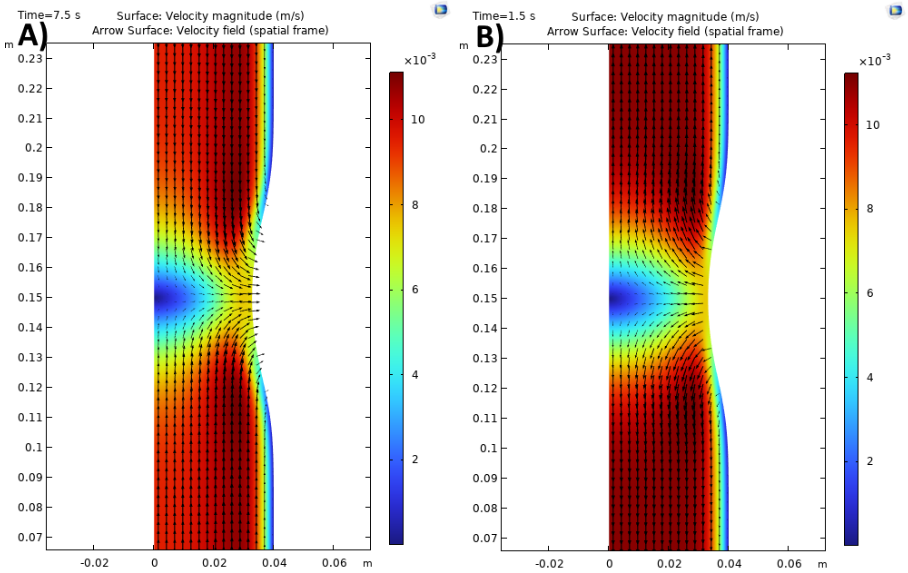


**Fig. 11**: A) Extensional flow B) Squeeze flow

From Fig. 11, the value of $L_t$ in the COMSOL simulation is approximately $0.1$ m. The liquid used in the COMSOL simulation has a density $\rho = 1000$ kg/m$^3$ and a viscosity $\mu = 0.01$ Pa.s. A higher viscosity liquid is chosen to ensure the liquid velocity profile remains laminar.

Since $U_{ar}$ value varies according to the hyperbolic tangent function or Eq. S.12, an average value for $U_{ar}$ must be used. To evaluate $U_{ar,avg}$, the following equation is used

$$U_{ar,avg} = \frac{U_{ar,max}}{L_t} \int_{0.1}^{0.2} \left[A_1 \tanh\left(A_2\left(Y - A_3\right)\right) - A_1\right] dY = 0.54 U_{max} \tag{S.14}$$

The average liquid velocity magnitude is evaluated at $Y = 0.25$ m as this is the region of interest since at this location, there is no effect on the geometry due to the deformation caused by contraction or relaxation.

Under the conditions $r_{fin} = 0.0307$ m at $Y = 0.15$ m, the location of the maximum contracted region, the COMSOL simulation reports an average liquid velocity magnitude of $0.00766$ m/s at $Y = 0.25$ m.

The average liquid velocity magnitude reported by the COMSOL simulation is compared using S.7. However, Eq. S.7 needs further derivation to evaluate the average liquid velocity magnitude, denoted as $u_{ae}$

$$u_{a,avg} = \frac{u_{a,max}}{2} = \frac{U_{ar,avg} L_t}{r_{fin}} \tag{S.15}$$

$$u_{ae} = u_{a,avg}\frac{r_{fin}^2}{r_{ini}^2} = \frac{U_{ar,avg}L_t r_{fin}}{r_{ini}^2} \tag{S.16}$$

For, $L_t = 0.1$ m, $r_{fin} = 0.0307$ m, $U_{ar,avg} = 0.003816$ m/s, $r_{ini} = 0.04$ m, conditions similar to those used in the COMSOL simulation, the average liquid velocity magnitude for the squeeze/extensional flow, calculated using Eq. S.16 is $0.00732$ m/s.

The value is quite comparable to that reported by the COMSOL simulation, thereby validating the derived equation.

##### 7.1.1 Extensional flow in the compartmental modeling framework

In our previous work [7], the authors derived equations that define a stretch-contraction transformation. These equations are expressed as follows

$$\omega_1 = \frac{L_d}{4} \tag{S.17}$$

$$\omega_d = \frac{\omega_1 - \frac{L_t}{4}}{2} \tag{S.18}$$

$$\omega_2 = \sqrt{\omega_1^2 - \omega_d^2} \tag{S.19}$$

$$\omega_3 = 2\omega_2 \tag{S.20}$$

In the model, the amplitude or maximum value of the deformed radius of the compartment was defined as $r_d$. Using the mean value theorem, the average value of the deformed radius $r_{d,avg}$ for a compartment is denoted as

$$r_{d,avg} = \frac{1}{L_t}\int_0^{L_t} r_d\, dL_t = \frac{1}{L_t}\left(\omega_3\omega_d + \frac{L_t r_d}{2}\right) \tag{S.21}$$

Since $r_d$ is the maximum value of the deformed radius and $U_{ar}$ in Eq. S.11 is derived based on $r_d$, $U_{ar}$ corresponds to the maximum contraction velocity. To account for the average contraction/relaxation velocity in a compartment, the expression for $U_{ar,avg}$ is denoted as:

$$U_{ar,avg} = \frac{r_{d,avg}U_{ar}}{r_d} \tag{S.22}$$

Inserting Eq. S.21 in Eq. S.22 and expressing in terms of $L_t$, $L_d$, $r_d$ and $U_{ar}$, $U_{ar,avg}$ is represented as

$$U_{ar,avg} = \frac{\left[\frac{L_t\, r_d}{2} + 2\sqrt{\frac{L_d{}^2}{16} - \left(\frac{L_d}{8} - \frac{L_t}{8}\right)^2}\,\left(\frac{L_d}{8} - \frac{L_t}{8}\right)\right] U_{ar}}{L_t\, r_d} \tag{S.23}$$

Using Eq. S.16 and Eq. S.23, the fluid velocity for the extensional/squeeze flow $u_{ae}$ in the compartmental modeling framework is denoted as

$$u_{ae} = \frac{\left[\frac{L_t\, r_d}{2} + 2\sqrt{\frac{L_d{}^2}{16} - \left(\frac{L_d}{8} - \frac{L_t}{8}\right)^2}\,\left(\frac{L_d}{8} - \frac{L_t}{8}\right)\right] U_{ar}\, r_{fin}}{r_d\, r_{ini}^2} \tag{S.24}$$

The volumetric flow rate for the extensional/squeeze flow $Q_{ae}$ in the compartmental modeling framework is denoted as

$$Q_{ae} = u_{ae}\,\pi\,r_{ini}^2 \tag{S.25}$$

#### 7.2 Compartmental model setup

Peristaltic behavior is a continuous process where the peristaltic wave propagates from the Pacemaker region (PR) to the TA of the stomach. To model this continuous behavior of gastric liquid caused by the peristaltic wave, the compartments are set up to preserve this continuity by enabling interaction between them. This interaction is modeled using differential equations that maintain the maximum velocity or flow rate of gastric liquid once each compartment reaches its maximum contraction during peristalsis. The maximum velocity or flow rate of gastric liquid during peristalsis is affected only when the subsequent compartment experiences a stronger contraction. The continuous behavior between compartments ceases at the TA, where the peristaltic wave dissipates, as demonstrated in a study by Ishida et al., 2019 [10].

To begin deriving the compartmental model interaction to predict gastric liquid behavior during peristalsis, the maximum peristaltic gastric liquid velocity is calculated in the PA and MA compartments for a given stimulus response $\kappa$.

Initially, the maximum membrane potentials of the ICC ($V_{m,ICC,max}$) and SMC ($V_{m,SMC,max}$) are calculated by solving the steady-state equations for Eq. 2 and Eq. 5. Using these steady-state equations, $V_{m,ICC,max}$ and $V_{m,SMC,max}$ are determined as

$$V_{m,ICC,max} = V_{mrest,ICC} + \kappa e^{-\left(s_{ICC}t_{cyc,peak}\right)^2} \tag{S.26}$$

$$V_{m,SMC,max} = \frac{G_{coup}V_{m,ICC,max} + V_{mrest,SMC}}{1 + G_{coup}e^{-\left(s_{ICC}t_{cyc,peak}\right)^2}} \tag{S.27}$$

After determining the maximum membrane potential of the SMC $V_{m,SMC,max}$, for a given stimulation value, $\kappa$, it is essential to compute the maximum intracellular calcium concentration and the maximum active stress in the muscle tissue. However, solving for these quantities directly using the mechanistic model equations, such as Eq. 6 and Eq. 7, is complex due to the intricate relationship between the SMC membrane potential and intracellular calcium concentration.

To overcome this challenge, polynomial curves are fitted to relate the SMC membrane potential to the number of cross bridges that influence the maximum maximum active stress in the muscle tissue. These fitted curves are represented by Eq. S.28 and Eq. S.29, respectively, and provide an easier method for calculating the desired quantities. The fitted curves are visualized in Fig. 12, providing a graphical representation of these relationships.

The actual curves used for fitting are derived from the mechanistic model equations outlined in the paper. This approach simplifies the calculation process and allows for more straightforward determination of the quantities influencing active tissue stress in response to the SMC membrane potential.

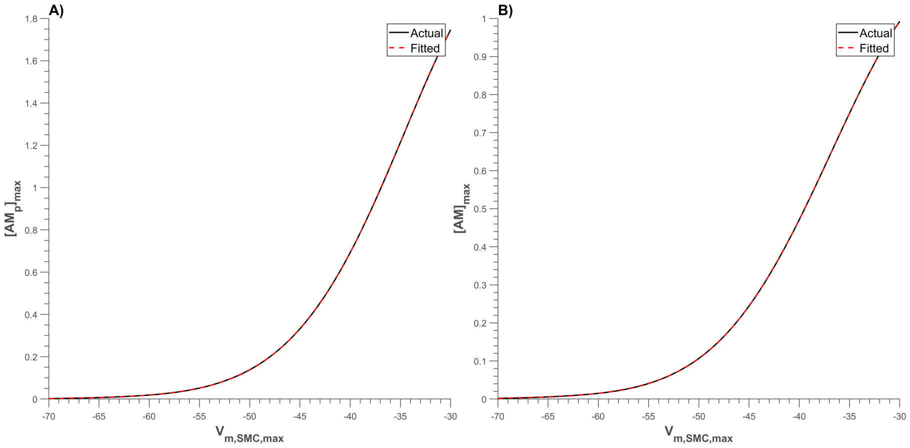


**Fig. 12**: Curve fitting that relates $V_{m,max,SMC}$ to the maximum formulation of cross-bridges in the muscle tissue A) $V_{m,max,SMC}$ to maximum concentration of actin phosphorylated myosin $[AM_p]_{max}$ B) $V_{m,max,SMC}$ to maximum concentration of actin myosin $[AM]_{max}$

$$[AM]_{max} = \sum_{z=1}^{7} \mathcal{D}_{i-1} V^{i}_{SMC,max} \tag{S.28}$$

$$[AM_p]_{max} = \sum_{z=1}^{6} \mathcal{E}_{i-1} V^{i}_{SMC,max} \tag{S.29}$$

Here $\mathcal{D}$ and $\mathcal{E}$ are the polynomial coefficients of the fitted curves. Using Eq. 7, the maximum active stress $\sigma_{c,max}$ in the muscle tissue is computed.

$$\sigma_{c,max} = \sigma_{max} \left( [AM_p]_{max} + [AM]_{max} \right) \tag{S.30}$$

To relate $\sigma_{c,max}$ to the maximum stretch in the muscle tissue $\lambda_{max}$, a curve is fitted using a natural log equation, represented as Eq. S.31. The actual curve computes the steady-state tissue stretch response for a given tissue stress. The comparison between the actual and the fitted curves for $\sigma_{c,max}$ and $\lambda_{max}$ is illustrated in Fig. 13.

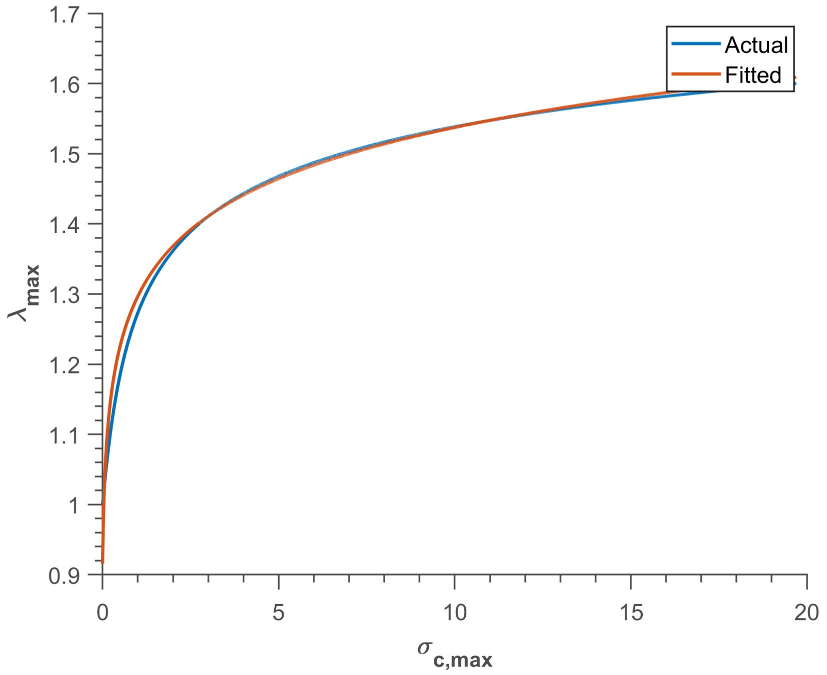


**Fig. 13**: Curve fitting that relates $\sigma_{c,max}$ to $\lambda_{max}$

$$\lambda_{max} = \mathcal{F}_1 \left( \ln \left( \left[ \mathcal{F}_2 \left( \sigma_{c,max} + \mathcal{F}_3 \right) \right]^{\mathcal{F}_4} \right) \right) + \mathcal{F}_5 \tag{S.31}$$

Here $\mathcal{F}$ is the coefficient of the fitted curve. Using $\lambda_{max}$, the maximum gastric liquid velocity and volumetric flow rate are calculated using algebraic equations from Eq. 12 to Eq. 26 in both scenarios: with the pyloric sphincter PS open and closed.

Since peristalsis only occurs in the PA, MA, and TA, the differential equations that describe the continuity of peristalsis are defined exclusively for these compartments.

For an open PS, the continuous volumetric flow rate of gastric liquid contents within a compartment is denoted as $Q_{pr,f*_i}$. To formulate the differential equation for $Q_{pr,f*_i}$, specific conditions for each compartment must be established individually.

For the PA compartment, the differential equation for $Q_{pr,f*_1}$ is set up such that when the peristaltic flow rate $Q_{pr,f_1}$ increases in the PA compartment, the value of $Q_{pr,f*_1}$ closely follows the path of $Q_{pr,f_1}$. This relationship is expressed by the differential equation $\frac{dQ_{pr,f*_1}}{dt} = k_{pr} \left( Q_{pr,f_1} - Q_{pr,f*_1} \right)$

where $k_{pr}$ is the rate at which $Q_{pr,f*_1}$ follows $Q_{pr,f_1}$. The value of $k_{pr}$ must be high so that $Q_{pr,f*_1}$ value can closely track $Q_{pr,f_1}$ value. For practical purposes, $k_{pr}$ is set to $10\,\mathrm{s}^{-1}$, which is high enough for $Q_{pr,f*_1}$ to follow $Q_{pr,f_1}$ value accurately while allowing efficient computation with an Euler solver.

Once $Q_{pr,f*_1}$ reaches the maximum flow rate value for the PA compartment, denoted as $Q_{pr,f*,max_1}$, this maximum value is maintained until the flow rate in the MA compartment, $Q_{pr,f_2}$, exceeds $Q_{pr,f*,max_1}$. When $Q_{pr,f_2}$ is greater than $Q_{pr,f*,max_1}$, the differential equation is $\frac{dQ_{pr,f*_1}}{dt} = -k_{pr}Q_{pr,f*_1}$. This results in $Q_{pr,f*_1}$ decreasing to 0 when $Q_{pr,f_2}$ exceeds $Q_{pr,f*,max_1}$ value indicating that the continuous peristaltic flow behavior now belongs to the MA compartment region.

The differential equation for the PA compartment, considering the conditions discussed above, is

$$\frac{dQ_{pr,f*_1}}{dt} = \begin{cases} k_{pr}\left(Q_{pr,f_1} - Q_{pr,f*_1}\right), & \text{if } Q_{pr,f*_1} < Q_{pr,f*,max_1} \\ & \text{and } Q_{pr,f_2} < Q_{pr,f*,max_1} \\ -k_{pr}Q_{pr,f*_1}, & \text{if } Q_{pr,f_2} > Q_{pr,f*,max_1} \\ 0, & \text{otherwise} \end{cases} \tag{S.32}$$

For the MA compartment, the value of the continuous flow rate of peristaltic behavior, $Q_{pr,f*_2}$, increases and follows the path of $Q_{pr,f_2}$ when $Q_{pr,f_2}$ is greater than $Q_{pr,f*,max_1}$. This value is maintained when $Q_{pr,f_2}$ reaches $Q_{pr,f*,max_2}$. The value of $Q_{pr,f*_2}$ then decreases to 0 when $Q_{pr,f_3}$ is greater than $Q_{pr,f*,max_2}$ indicating that the continuous peristaltic flow behavior now belongs to the TA compartment region.

The differential equation for the MA compartment, considering the conditions discussed above, is represented as

$$\frac{dQ_{pr,f*_2}}{dt} = \begin{cases} & \text{if } Q_{pr,f_2} > Q_{pr,f*,max_1} \\ k_{pr}\left(Q_{pr,f_2} - Q_{pr,f*_2}\right), & \text{and } Q_{pr,f*_2} < Q_{pr,f*,max_2} \\ & \text{and } Q_{pr,f_3} < Q_{pr,f*,max_2} \\ -k_{pr}Q_{pr,f*_2}, & \text{if } Q_{pr,f_3} > Q_{pr,f*,max_2} \\ 0, & \text{otherwise} \end{cases} \tag{S.33}$$

For the TA compartment, the value of the continuous flow rate of peristaltic behavior, $Q_{pr,f*_3}$, increases and follows the path of $Q_{pr,f_3}$ when $Q_{pr,f_3}$ is greater than $Q_{pr,f,max_2}$. Since the TA is the region where the peristaltic contraction ends, the volumetric flow rate influenced by peristalsis drops to 0 when the TA peristaltic contraction relaxes $(U_{sq} < 0)$.

The differential equation for the TA compartment, considering the conditions discussed above, is denoted as

$$\frac{dQ_{pr,f*_3}}{dt} = \begin{cases} k_{pr}\left(Q_{pr,f_3} - Q_{pr,f*_3}\right), & \text{if } Q_{pr,f_3} > Q_{pr,f*,max_2} \\ -k_{pr}Q_{pr,f*_3}, & \text{if } U_{sq} < 0 \\ 0, & \text{otherwise} \end{cases} \tag{S.34}$$

When the PS is closed, a retrograde flow occurs. The peristaltic velocity in each compartment when the PS is closed is calculated using a similar framework as that for the forward flow. To simplify calculations and maintain a consistent framework, the direction is ignored for both forward and reverse flow by taking the absolute value of the flow rate or velocity. Once the calculations are completed, the direction is assigned using a negative sign for gastric emptying flow and a positive sign for retrograde flow (gastric mixing). The differential equations that represent the continuous peristaltic flow in each compartment when the PS is closed, $u_{pr,b*_i}$ where $(i = 1, 2, 3)$, are denoted as follows

$$\frac{du_{pr,b*_1}}{dt} = \begin{cases} k_{pr}\left(u_{pr,b_1} - u_{pr,b*_1}\right), & \text{if } u_{pr,b*_1} < u_{pr,b*,max_1} \\ & \text{and } u_{pr,b_2} < u_{pr,b*,max_1} \\ -k_{pr}u_{pr,b*_1}, & \text{if } u_{pr,b_2} > u_{pr,b*,max_1} \\ 0, & \text{otherwise} \end{cases} \tag{S.35}$$

$$\frac{du_{pr,b*_2}}{dt} = \begin{cases} k_{pr}\left(u_{pr,b_2} - u_{pr,b*_2}\right), & \text{if } u_{pr,b_2} > u_{pr,b*,max_1} \\ & \text{and } u_{pr,b*_2} < u_{pr,b*,max_2} \\ & \text{and } u_{pr,b_3} < u_{pr,b*,max_2} \\ -k_{pr}u_{pr,b*_2}, & \text{if } u_{pr,b_3} > u_{pr,b*,max_2} \\ 0, & \text{otherwise} \end{cases} \tag{S.36}$$

$$\frac{du_{pr,b*_3}}{dt} = \begin{cases} k_{pr}\left(u_{pr,b_3} - u_{pr,b*_3}\right), & \text{if } u_{pr,b_3} > u_{pr,b*,max_2} \\ -k_{pr}u_{pr,b*_3}, & \text{if } U_{sq} < 0 \\ 0, & \text{otherwise} \end{cases} \tag{S.37}$$

##### 7.3 PS contraction

The contraction period of the PS is longer than that of the antrum, as demonstrated in both an experimental study by Indireshkumar et al., 2000 [47] and a computational study by Ishida et al., 2019 [10]. However, the cause of this longer contraction period in the PS has not been addressed in the literature. Therefore, this paper proposes two models to explain why the PS contraction period is longer compared to the antrum.

The first model proposes a change in the soft tissue material properties of the PS. During the relaxation of PS tissue, it follows a path that results in a wider hysteresis loop compared to the antrum, as shown in Fig. 15. This characteristic allows the PS to have a longer contraction period without altering the width of the ICC “slow waves," as shown in Fig. 14 B). The parameters accounting for a wider hysteresis curve at the PS are illustrated in Table 4.

The second model suggests that the phasic ICC “slow waves" are wider in the PS compared to the antrum, as illustrated in Fig. 14 A). The parameters for the ICC “slow waves" and muscle tissue are presented in Table 3. These parameter values are similar to those reported in a previous study by Fernandes et al., 2024 [7], with a few modifications to account for the wider ICC “slow waves" that result in the longer contraction period of the PS.

**Table 3**: Parameters for model 1

| Parameter | Value | Unit |
|---|---|---|
| $t_{open_{1/2/3/4}}$ | 7 | s |
| $S_{1/2/3/4/5/6_{1/2/3}}$ | 0.45 / 0.89 / 4.2 / 0.19 / 2 / 5 | kPa / – / kPa·s / kPa$^{-1}$ / – / – |
| $S_{1/2/3/4/5/6_4}$ | 0.45 / 0.89 / 4.2 / 0.22 / 2 / 20 | kPa / – / kPa·s / kPa$^{-1}$ / – / – |

Note: Tissue constants $S_{1/2/3/4/5/6}$ are listed in sequence. The dash (–) indicates dimensionless entries.

**Table 4**: Parameters for model 2

| Parameter | Value | Unit |
|---|---|---|
| $t_{open_{1/2/3}}$ | 7 | s |
| $t_{open_4}$ | 9 | s |
| $S_{1/2/3/4/5/6_{1/2/3/4}}$ | 0.45 / 0.89 / 4.2 / 0.19 / 2 / 5 | kPa / – / kPa·s / kPa$^{-1}$ / – / – |

Note: Same convention as Table 3 for the constants $S_{1/2/3/4/5/6}$.

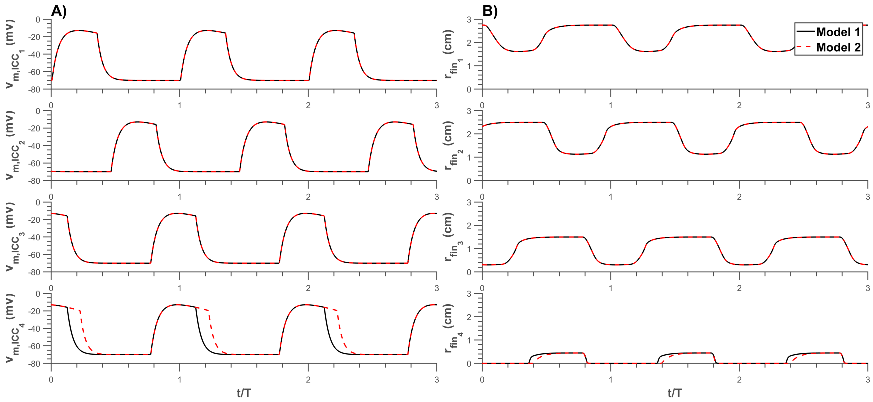


**Fig. 14**: Model 1 and model 2 behaviors for A) Membrane potential of ICC “slow waves and B) Radius of compartments

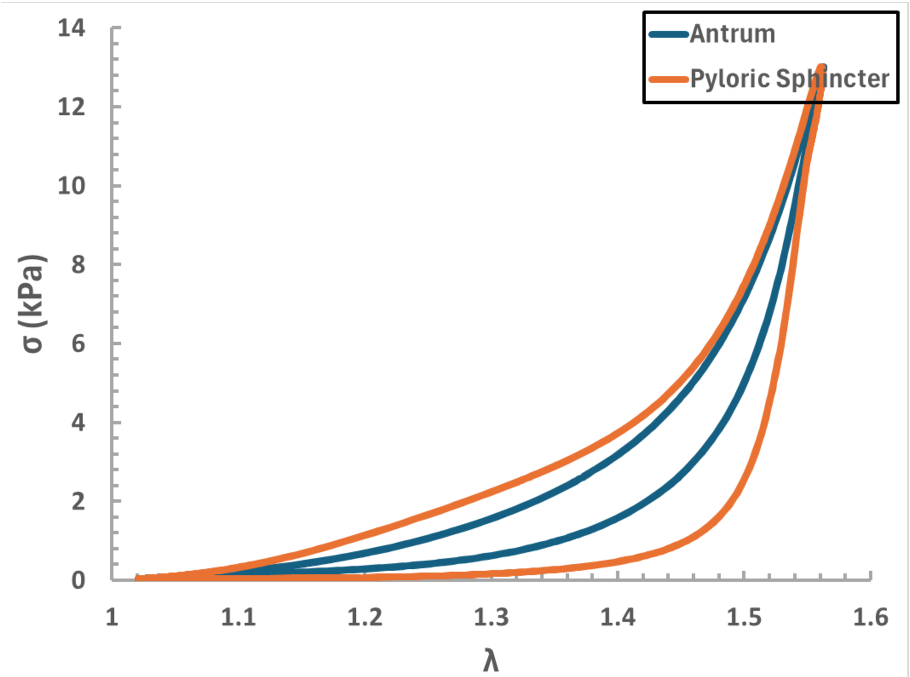


**Fig. 15**: Soft tissue material model behavior for the antrum and PS compartment in model 1

In Fig. 15, the parameters were adjusted to achieve a wider hysteresis loop for the PS, resulting in a faster response rate and longer contraction period compared to the antrum of the stomach. These parameters, which produce the stress versus stretch curve shown in Fig. 15, are presented in Table 3.

In this study, model 1 is preferred over model 2 for computing the results because, as shown in Fig. 14 B), the PS compartment relaxes quicker in model 1 than in model 2. This behavior is desirable as it aligns with findings from previous studies in the literature [10, 47].

#### 7.4 Parameters

##### 7.4.1 Healthy stomach

The parameters in this study for a healthy human stomach can be found in Table 5

**Table 5**: Parameters for human healthy stomach

| Parameter | Value | Unit | Reference |
|---|---|---|---|
| | Electrical | | |
| $c_{1/2/3}$ | 0.25 / 0.25 / 0.6 | cm/s | [7, 10] |
| $\kappa_{1/2/3/4}$ | 59.53 | mA | [7] |
| $V_{m,rest,ICC}$ | -70 | mV | [2] |
| $V_{m,rest,SMC}$ | -70 | mV | [1] |
| $s_{ICC}$ | 0.05 | $s^{-1}$ | [7] |
| $R_{ICC}$ | 1 | Ω | [7] |
| $\mathcal{C}_{ICC}$ | 1 | F | [7] |
| $s_{SMC}$ | 0.05 | $s^{-1}$ | [7] |
| $R_{SMC}$ | 1 | Ω | [7] |
| $\mathcal{C}_{SMC}$ | 1 | F | [7] |
| $t_{open}$ | 7 | s | [7] and Section 7.3 |
| $t_{end_{1/2/3/4}}$ | 20 | s | [7] |
| $G_{coup_{1/2/3/4}}$ | 0.63 / 1.26 / 1.26 / 1.26 | S | [7, 80] |
| | Mechanical | | |
| $\sigma_{max}$ | 238.3 | kPa | [7] |
| $S_{1/2/3/4/5/6_{1/2/3}}$ | 0.45 / 0.89 / 4.2 / 0.19 / 2 / 5 | kPa / – / kPa·s / $kPa^{-1}$ / – / – | [7] |
| $S_{1/2/3/4/5/6_4}$ | 0.45 / 0.89 / 4.2 / 0.22 / 2 / 20 | kPa / – / kPa·s / $kPa^{-1}$ / – / – | [7] and Section 7.3 |
| $Lt_{1/2/3/4}$ | 2.4 / 2.4 / 2.1 / 6 | cm | [7, 10] |
| | Fluid | | |
| $D_*$ | 2.5 | cm | [7] |
| $K$ | 1.5 | – | [7] |
| $O_{1/2}$ | 0.05 / 0.92 | – | [7] |
| $k_{pr}$ | 10 | $s^{-1}$ | [7] |

Note: Dashes (–) indicate dimensionless parameters.

##### 7.4.2 Parameters for the stomach with altered “slow wave" frequency and amplitude

The parameters modified to simulate impaired coordination of the “slow wave" frequency between the antrum and the PS are listed in Tables 6 and 7. Parameters adjusted to model coordinated antrum–PS activity under bradygastric and tachygastric conditions are provided in Table 8. Finally, the parameters used to represent quiescent “slow wave" activity are shown in Table 9.

**Table 6**: Change in PS “slow wave" frequency parameter

| Parameter | Value | Unit |
|---|---|---|
| $t_{end_{1/2/3}}$ | 20 | s |
| $t_{end_4}$ | [10,60] | s |

Range indicates parameter sweep used in simulations.

**Table 7**: Change in antrum “slow wave" frequency parameter

| Parameter | Value | Unit |
|---|---|---|
| $t_{end_{1/2/3}}$ | [10,60] | s |
| $t_{end_4}$ | 20 | s |

Range indicates parameter sweep used in simulations.

**Table 8**: Change in stomach (antrum + PS) “slow wave" frequency parameter

| Parameter | Value | Unit |
|---|---|---|
| $t_{end_{1/2/3/4}}$ | [10,50] | s |

Range indicates parameter sweep used in simulations.

**Table 9**: Change in “slow wave" amplitude parameter

| Parameter | Value | Unit |
|---|---|---|
| $\kappa_{1/2/3}$ | $\kappa_*$ | mA |
| $\kappa_{end_4}$ | $\kappa$ | mA |

$\kappa = 59.53$ mA; $\kappa_*$ can refer to $\kappa_A$, $\kappa_B$, or $\kappa$.
